# Molecular dynamics reveals complex compensatory effects of ionic strength on the SARS-CoV-2 Spike/hACE-2 interaction

**DOI:** 10.1101/2020.08.25.267351

**Authors:** Anacleto Silva de Souza, Jose David Rivera, Vitor Medeiros Almeida, Pingju Ge, Robson Francisco de Souza, Chuck Shaker Farah, Henning Ulrich, Sandro Roberto Marana, Roberto Kopke Salinas, Cristiane Rodrigues Guzzo

**Affiliations:** Department of Microbiology, Institute of Biomedical Sciences, University of São Paulo, São Paulo, Brazil; Department of Biochemistry, Institute of Chemistry, University of São Paulo, São Paulo, Brazil; Acrobiosystems Inc., Beijing 100176, China

**Keywords:** COVID-19, pandemic, coronavirus, spike, ACE-2

## Abstract

The SARS-CoV-2 pandemic has already killed more than 800,000 people worldwide. To gain entry, the virus uses its spike protein to bind to host hACE-2 receptors on the host cell surface and mediate fusion between viral and cell membranes. As initial steps leading to virus entry involves significant changes in protein conformation as well as in the electrostatic environment in the vicinity of the spike-hACE-2 complex, we explored the sensitivity of the interaction to changes in ionic strength through computational simulations and surface plasmon resonance. We identified two regions in the receptor-binding domain (RBD), E1 and E2, which interact differently with hACE-2. At high salt concentration, E2-mediated interactions are weakened but are compensated by strengthening E1-mediated hydrophobic interactions. These results provide a detailed molecular understanding of spike RBD/hACE-2 complex formation and stability under a wide range of ionic strengths.

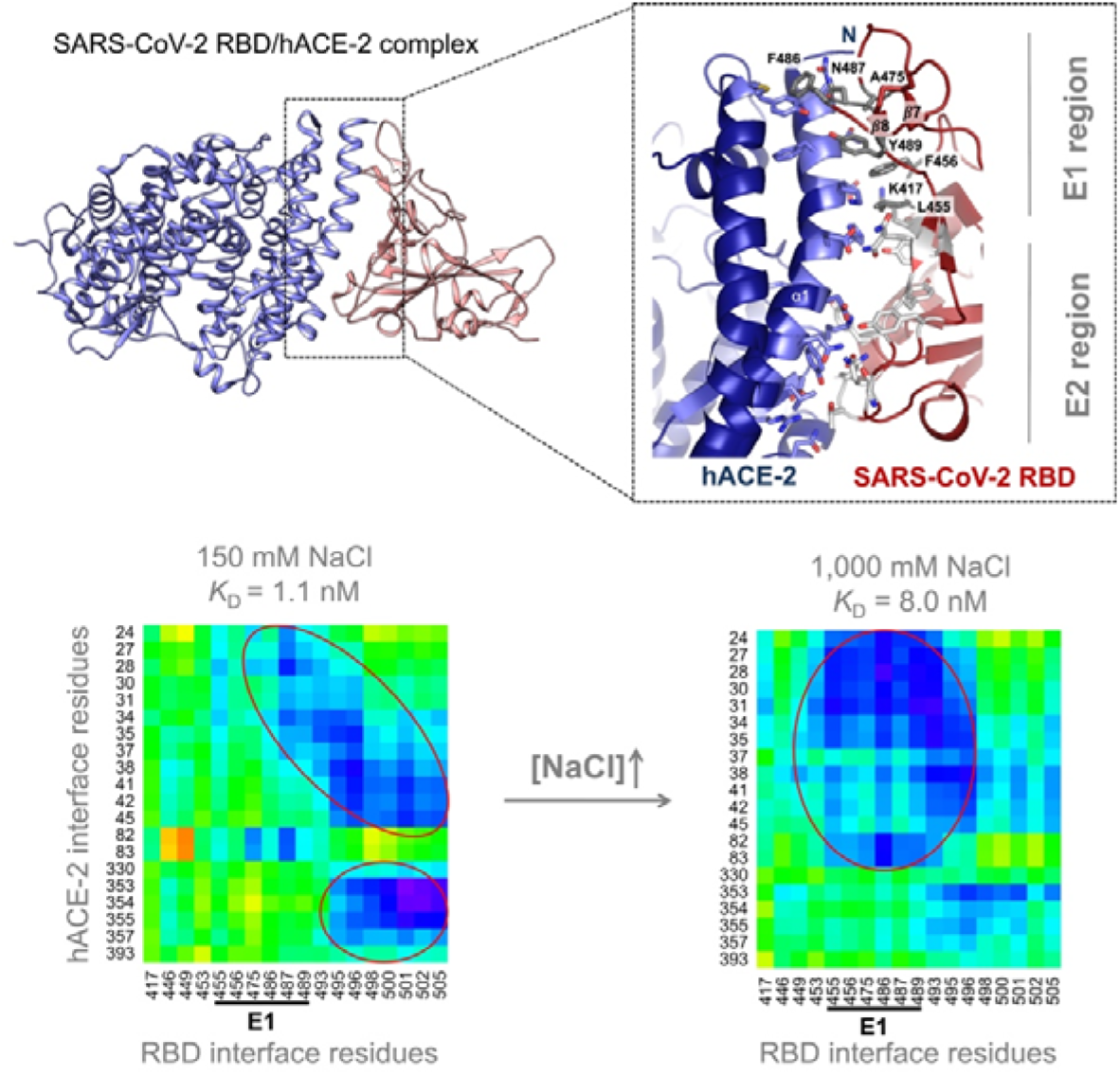

---

Severe Acute Respiratory Syndrome Coronavirus 2 (SARS-CoV-2) is the etiological agent of the current outbreak of coronavirus disease (COVID-19). From late 2019 and into August 2020, SARS-CoV-2 has already infected more than 22 million and killed more than 800,000 people in 216 countries ^1^. The cell entry mechanism is thought to be initiated by the binding of the viral Spike glycoprotein to the human angiotensin-converting enzyme (hACE-2) ^2,3^ expressed on the surface of airway epithelial ^4^, kidney, intestine, macrophages and other cells ^5^. Spike is a membrane surface trimeric protein, which contains active S1 and S2 subunits. The S1 subunit is involved in host cell receptor-binding and S2 mediates cell and viral membrane fusion ^3,6,7^. The hACE-2 receptor consists of C-terminal collectrin-like and N-terminal peptidase domains (NPD), responsible for the enzymatic activity for cleaving angiotensin II to angiotensin (1-7). The Spike S1 subunit contains a highly variable receptor-binding domain (RBD) comprising residues 331-524 (**Figure S1**) ^3,6,7,8^. Binding of the Spike-RBD to hACE-2 NPD triggers a conformational change exposing spike S1/S2 and the S2’ sites for cleavage by furin and transmembrane protease serine 2 (TMPRSS2)^9^, activating membrane fusion and hence virus entry ^10–13^.

Structural studies showed that each Spike RBD interacts with one hACE-2 NPD domain through hydrogen bonding with few salt bridges and hydrophobic contacts^14,15^. The spike RBD and hACE-2 complex (RBD/hACE-2) has a high binding affinity with dissociation constants (*K*_D_) ranging from 5.1 to 44.2 nM ^15,16^. During membrane fusion, Spike undergoes significant conformational changes in an environment whose electrostatic nature is also highly variable, both temporally and spatially ^17^ and so the RBD-hACE-2 interaction must remain stable during membrane approximation, proteolytic cleavage at S2’ and dissociation of the S1 and S2 subunits prior to S2-mediated membrane fusion. During this process, it is reasonable to envision significant fluctuations in the electrostatic environment in the vicinity of the RBD/hACE-2 complex. We therefore investigated how ionic strength affects complex stability using computational simulations and SPR experiments.

We characterized the molecular properties of the SARS-CoV-2 RBD/hACE-2 interface by mapping hydrophilic and hydrophobic profiles for each amino acid residue (Figure 1a and **Table S1**). The main differences between amino acid sequences of SARS-CoV and SARS-CoV-2 Spike proteins are located on the RBD interface with hACE-2 (Figure 1a-c). These differences alter the hydropathy profile of the RBD residues located at the interface (Figures 1c and 1d). From this analysis, we named two regions, E1 and E2 (Figure 1). The E1 region corresponds to residues 417, 455-456, 470-490. The E1 region in SARS-CoV-2 is slightly more hydrophobic than the E1 region in SARS-CoV, especially due to substitution of Tyr to Leu at position 455 and Pro to Ala at position 475, which is partly compensated by the substitution of Val to Lys at position 417 (Figures 1e; **Table S1**). In contrast, the E2 region, made up of residues 444-454 and 493-505 (Figure 1e), which mediates mostly polar interactions with hACE-2, is slightly more hydrophilic in SARS-CoV-2 (Figures 1c and e; **Table S1**) mainly due to substitutions Tyr to Gln at position 498 and Thr to Asn at position 501 (all SARS-CoV-2 numbering; Figure 1e; **Table S1**).

**Figure 1.**
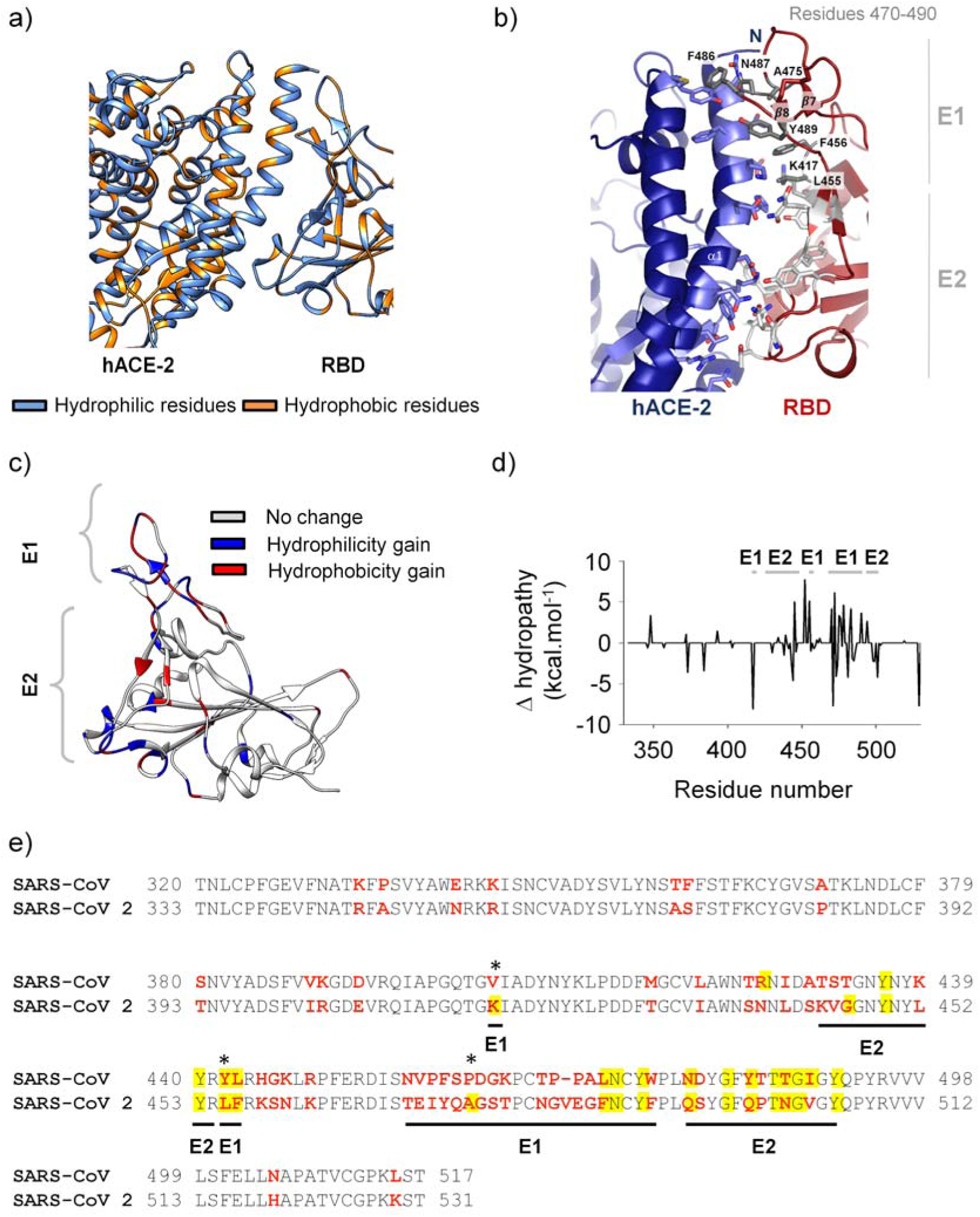
Properties of SARS-CoV-2 RBD interaction interfaces. Amino acid sequence and hydropathy analysis revealed two distinct regions at the SARS-Cov-2 RBD/hACE-2 interface. Based on structural analysis and hydropathy maps, the E1 region was delimited by K417 and 470-490 residues and by the hydrophobic cage formed by L455 and F456. The E2 region corresponded to 446-453 and 493-505 residues. (a) Hydropathy scale of SARS-CoV-2 RBD and hACE-2 color coded on the X-ray structure (PDB ID: 6M0J) (orange: hydrophobic; blue: hydrophilic). (b) RBD/hACE-2 interface residues in the E1 and the E2 regions. (c) SARS-CoV-2 RBD/hACE-2 (PDB ID: 6M0J) crystal structure colored according to the hydropathy scale differences between SARS-CoV-2 and SARS-CoV. (d) Hydropathy scale differences plotted as a function of the amino acid residue sequence. Positive values indicate a hydrophobicity gain in SARS-CoV-2 relative to SARS-CoV. (e) Multiple protein sequence alignment of SARS-CoV (GenBank: AAP41037.1) and SARS-CoV-2 (GenBank: QIG55994.1) Spike RBD. Residues that undergo significant change in hydropathy are indicated with asterisk.

In order to dissect the relative contribution of the E2 and E1 regions to the stable maintenance of the SARS-CoV-2 RBD/hACE-2 complex and the potential impact of the ionic environment in the binding affinity, we first sampled the conformational space of the complex (PDB ID 6M0J) using 100 ns of molecular dynamics (MD) at 150, 500, and 1,000 mM NaCl concentrations (Figure 2). Analysis of the backbone root-mean-square fluctuation (RMSF) showed that the RBD residues 470-490 (in E1 region) and hACE-2 residues 129-142 (distant from the interface) experienced the largest motions (Figure 2a and 2b). However, both were substantially reduced at higher ionic strengths (1,000 mM NaCl) (Figure 2a and 2b). Notably, hACE-2 sampled two major conformational states at 150 mM NaCl (state 1 of RMSD of ~2.4 Å and state 2 of ~3.4 Å) that are characterized by an opening-closing motion of residues 129-142, which is significantly affected by the salt concentration (Figure 2c and 2e, **Figure S2** and **S3**). Interestingly, probability distributions of the backbone RMSD along the MD trajectory showed that the RBD sampled a narrower conformational ensemble at higher salt (1,000 mM NaCl) in comparison to lower salt (150 mM NaCl) (Figure 2d). Most significantly the RBD/hACE-2 complex experienced fluctuations of the interface opening angle (Figure 2f and 2g). Average opening angles of 25°, 26°, 28° were observed at 150, 500 and 1,000 mM NaCl (Figure 2g and **Table S2**), respectively, also reflected in slightly greater RBD and hACE-2 center-of-mass distances (**Figure S4** and **Table S3**). This effect may be due to weakened polar contacts between hACE-2 and RBD at the E2 region. Indeed, interfacial hydrogen bonding analysis revealed that increased ionic strengths weakened some of the interactions between polar residue pairs in this region. Although some polar interactions in the E1 region are affected by increasing ionic strength such as in the case of N487^RBD^ that disrupts its interaction to Y83^hACE-2^ to perform intermolecular interactions. Some interactions are replaced to another as in the case of K417^RBD^ that decreases hydrogen bond (HB) occupancy to D30^hACE-2^ and interacts with H34^hACE-2^ and Q406^RBD^ at 1,000 mM NaCl when compared to 150 mM NaCl. Novel interactions in the E1 region were also observed in 1,000 mM NaCl as the case of interaction between Y489^RBD^ and T27^hACE-2^ and T83^hACE-2^ (**Table S4**). These compensatory effects together with the hydrophobic cage (L455 and F456) in the E1 region may allow the integrity of the complex in a broad range of salt concentrations. These observations are consistent with the nature of E1 and E2 regions.

**Figure 2.**
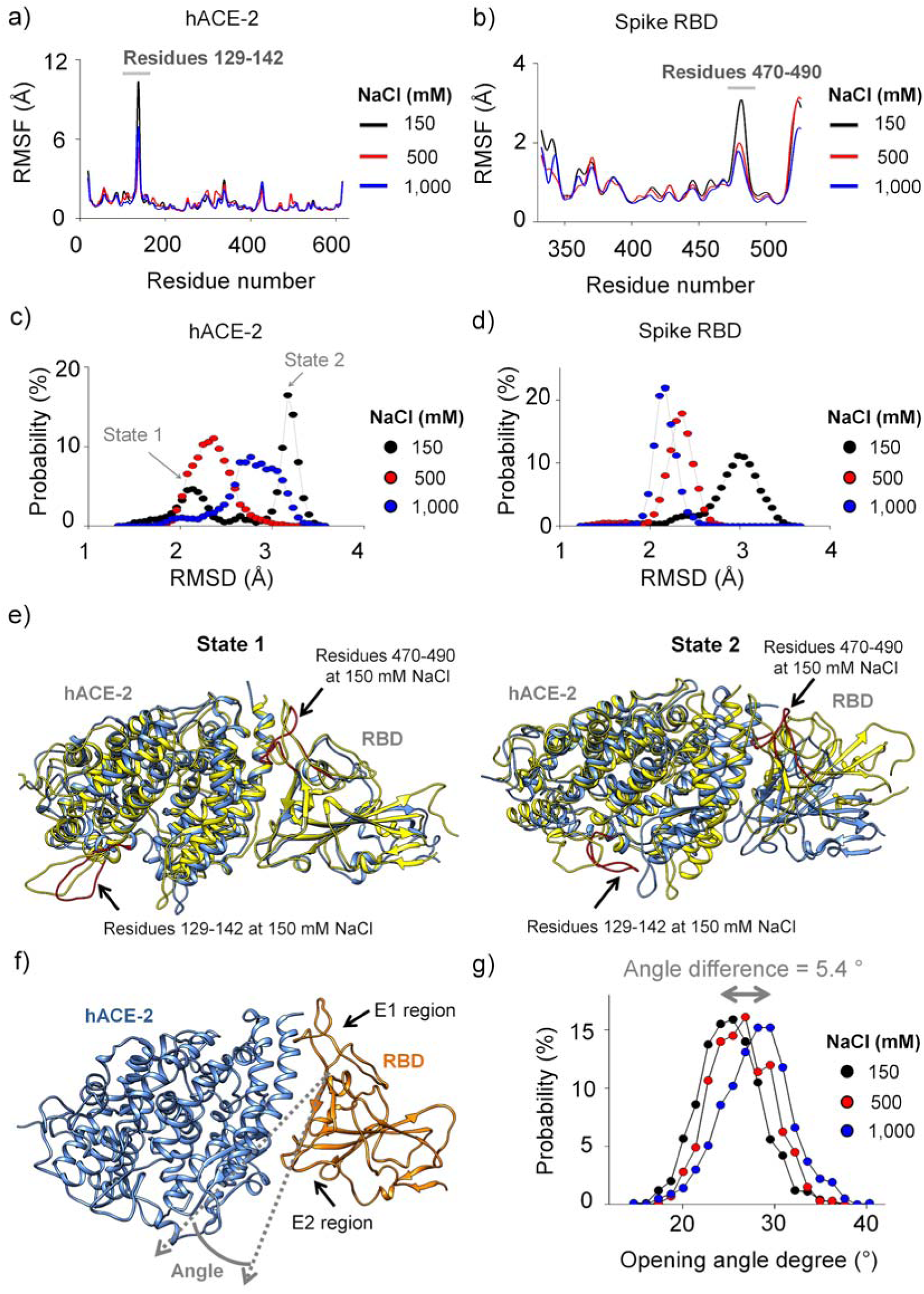
Structural analysis of the RBD/hACE-2 complex along the molecular dynamics simulations performed at increasing NaCl concentrations. Root-mean-square fluctuation (RMSF) of (a) hACE-2 and (b) RBD at different salt concentrations. (a) hACE-2 and (b) RBD RMSD probability distributions. (e) hACE-2 visits two conformational states in the presence of 150 mM NaCl (marked in blue) that differ mainly in the 129-142 residues (marked inthe red). The models were superimposed on a snapshot taken from the hACE-2/RBD trajectory performed at 500 mM NaCl (left; yellow) or 1,000 mM NaCl (right; yellow). RBD 470-490 residues are colored in red. (f) Opening of interaction interfaces as a function of the salt concentration. The opening is quantified based on the angle between the two vectors (gray dotted arrows) corresponding to the C_α_ coordinates of hACE-2 α-helix (residues 38-49) and Spike RBD β-strand (492-494 residues). (g) Probability distributions of the angles (panel f) at different salt concentrations. In all cases black refers to 150 mM, red to 500 mM, and blue to 1,000 mM NaCl.

We analyzed the covariance matrix of the spatial displacements of C atoms to assess internal motions of the RBD/h-ACE-2 complex (**Figure S6a**). This approach was based on the rationale that residues interacting through non-covalent bonds should, even afar, exert influence upon each other. A cut-off of 0.8 was adopted for the identification of significant positive and negative covariations. Covariant pairs were highly concentrated in secondary structure elements of the two proteins (**Figure S6b**). However, sets of covariant pairs changed at increasing NaCl concentrations. At 150 mM NaCl, we computed 3,558 covariant residue pairs for the spike RBD/hACE-2 complex. Surprisingly, this number decreased to 2,347 at 500 mM NaCl and increased back to 3,892 at 1,000 mM NaCl (Figure 3a). Most of the covariances were intramolecular, within the RBD or hACE-2, whereas only 18, 1 and 7 intermolecular covariances were observed at 150, 500 and 1,000 mM NaCl, respectively (Figure 3b). Among them a few negative covariances (11 and 3 at 150 and 1000 NaCl, respectively) were present (Figure 3b). While the increase of NaCl concentration caused similar changes in the number of intramolecular covariances, i.e., around 30% of them were disrupted at 500 mM NaCl and recovered at 1,000 mM NaCl, almost all intermolecular covariances (~ 95%) were disrupted at 500 mM NaCl and only 40% of them were re-established at 1,000 mM NaCl.

**Figure 3.**
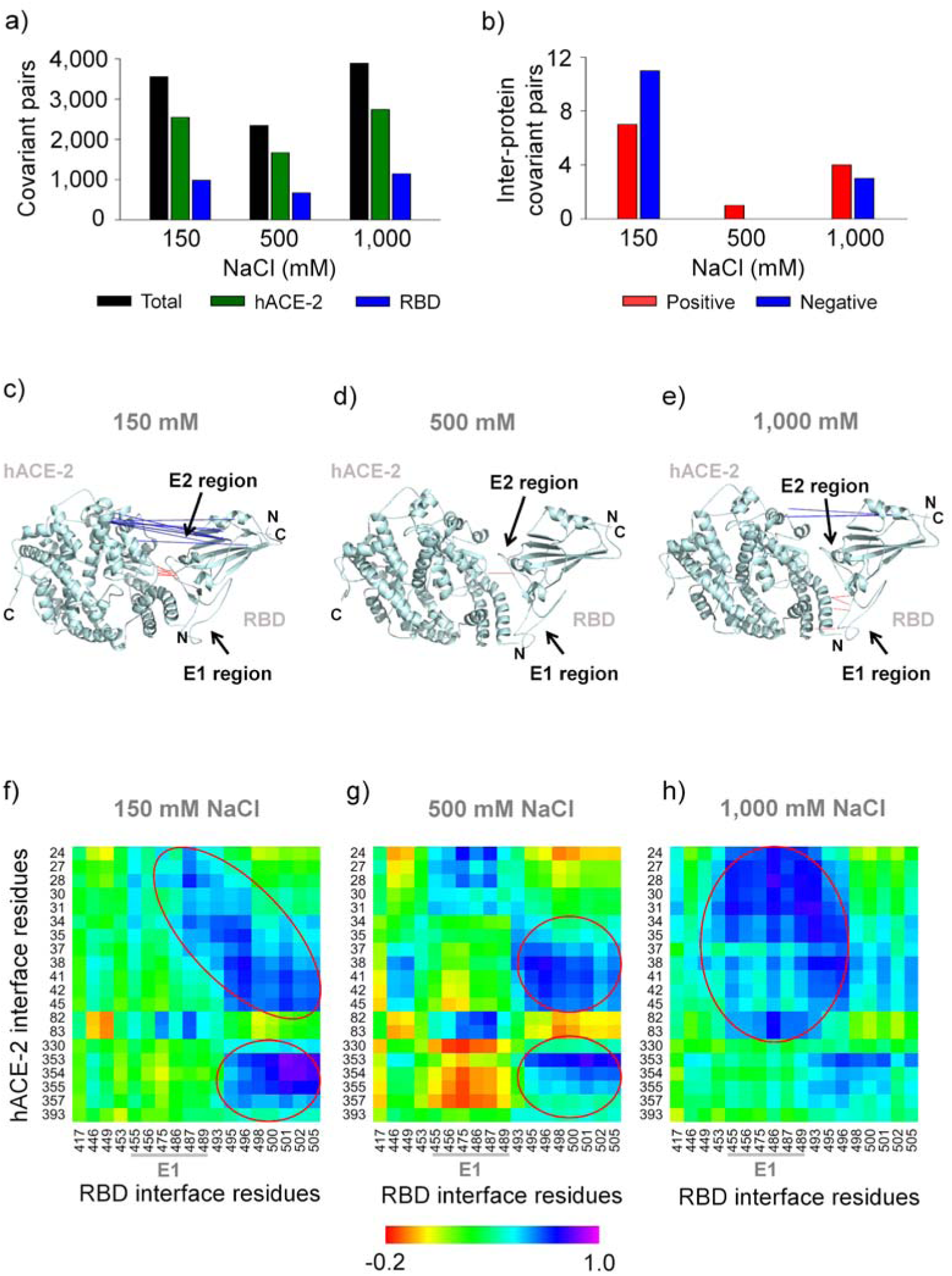
Covariance analysis of MD trajectories performed at increasing NaCl concentrations. (a) Total of covariant pairs observed in the RBD/hACE-2 complex at different salt concentrations. (b) Residues forming intermolecular covariant pairs mapped on the RBD/hACE2 complex at (c) 150 mM, (d) 500 mM, and (e) 1,000 mM NaCl. Residue pairs exhibiting positive covariance are connected by red lines, whereas negative covariant pairs are marked in blue. Displacement correlation matrix at the RBD/hACE-2 interface, with regions displaying positive covariances indicated by red ellipses: (f) 150 mM, (g) 500 mM, and (h) 1,000 mM NaCl. Pairs of C_α_ exhibiting covariance higher than 0.8 (positive and negative) were analyzed.

Residues forming intermolecular covariant pairs were identified (**Table S5**). Furthermore, the seven pairs of positively correlated residues at 150 mM NaCl are located in the hydrophilic E2 region at the interface of the RBD/hACE-2 complex. They were almost completely disrupted at 500 mM NaCl, and four new covariant pairs were formed at the opposite side of the complex interface at 1,000 mM NaCl solution (E1 region on the RBD) (Figure 3c-e). In order to better visualize the changes occurring as a function of the ionic strength, we also built a cross-correlation matrix of the complete set of residues, previously reported to belong to the complex interface using a cut-off of 4 Å^2^ (Figure 3f-h). Remarkably, RBD residues participating in the higher covariance pairs at 1,000 mM NaCl differed from those observed at 150 mM (Figure 3f-h; compare red ellipses). Therefore, ionic strength increments changed the interacting regions of the complex interface, suggesting that the RBD E1, rather than the E2 region, becomes more firmly attached to hACE-2 at higher ionic strengths (Figure 3c-h, **Table S5**).

We then investigated the relative contributions of the E1 and E2 regions to complex stability *in silico* by applying a harmonic force with constant velocity along the dissociation pathway (**Figure S5a, supplementary movies**). The complex (PDB ID 6M0J) was initially relaxed for 100 ns of MD in each NaCl concentration, and a pulling force was then applied on the RBD center-of-mass while the positions of the hACE-2 atoms were restricted. Along the dissociation trajectory, the intermolecular S19^hACE-2^/G476^RBD^ distance remained invariant for ~0.5 and 0.8 ns at 150 and 1,000 mM NaCl, respectively (**Figure S5b, supplementary movies**), suggesting that the intermolecular interaction at the E1 region became stronger at 1,000 mM NaCl. This effect may result from hydrophobic interactions. In the presence of 500 mM NaCl, the initial distance was around ~13 Å and was maintained for 0.4 ns, in contrast to the initial distance of 5-6 Å at 150 and 1,000 mM NaCl. The pulling force experiment indicated the presence of weaker intermolecular interactions at 500 mM NaCl compared to further NaCl concentrations, since the dissociation occurred earlier in the simulation at this ionic strength. This is consistent with the absence of intermolecular covariances at 500 mM NaCl. However, the T52^hACE-2^/G502^RBD^ distance at the E2 region remained invariant for ~ 0.5 ns at all salt concentrations (**Figure S5c, supplementary movies**).

To further characterize the roles of E2 and E1 regions in RBD/hACE-2 complex stability at the different salt concentration, we simulated RBD/hACE-2 reassociation beginning with the centers of masses of the two proteins separated by ~60 Å (Figure 4a-c). We observed that RBD readily associates with hACE-2 after ~2 ns and ~5 ns at 150 and 1,000 mM NaCl, respectively (Figures 4b and 4c), restoring the native interface (**Figure S7**). We noted that the RBD/hACE-2 center-of-mass separation at 1,000 mM is slightly greater than at 150 mM NaCl, as observed in the previous MD trajectory (**Figure S4** and **Table S3**). However, association was not observed at 500 mM NaCl when initiated at ~60 Å separation and could only be observed when beginning the simulation at ~30 Å (data not shown). This is consistent with the cross-correlation analysis that showed no intermolecular covariances at 500 mM NaCl (Figure 3d).

**Figure 4.**
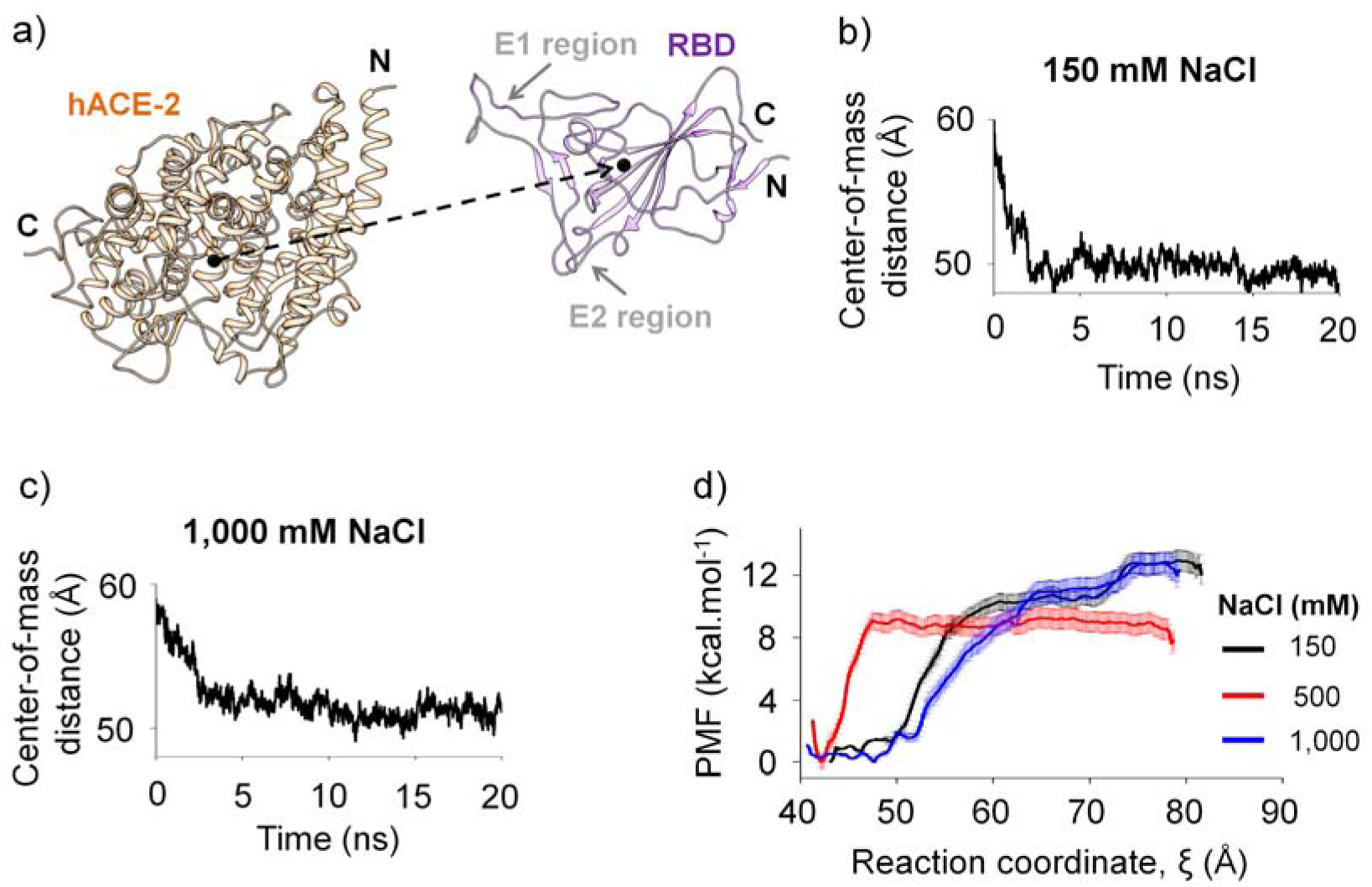
(a) - (c) RBD and hACE-2 reassociate after being separated by ~60 Å as indicated by the behavior of the center-of-mass distance as a function of the simulation time. (d) Potential of mean force (PMF) profiles obtained using the weighted histogram analysis method (WHAM) at different salt concentrations (150 mM, black; 500 mM, red; 1,000 mM, blue). The uncertainties were computed using 100 rounds of bootstrapping analysis. The reaction coordinate (ξ) was defined as the average intermolecular vector at the interface. The structure was rotated such that the Z-axis became parallel to the reaction coordinate vector. The value of ξ corresponds to the Z-component of the center-of-mass distance projected on the reaction coordinate. The energy reference level was set to the minimum corresponding to ξ = 43.1, 42.2, and 47.7 Å at 150, 500, and 1,000 mM NaCl, respectively.

Distance profiles were fitted to an exponential equation at 150 and 1,000 mM NaCl, each representing a free diffusion kinetics (Figure 4b-c and **Table S6**). Association rates, *k*, of 1.18 and 0.47 ns^-1^ were obtained at 150 and 1,000 mM NaCl, respectively (**Table S6**). The slower association rate observed at 1,000 mM NaCl could be due to weaker electrostatic interactions at higher salt concentration. Indeed, a greater number of Na^+^ or Cl^−^ ions is observed in the vicinity of the amino acids of the interface at 1,000 mM relative to 150 mM NaCl, particularly near charged residues mainly in hACE-2 and in the RBD E2 region (**Tables S7** and **S8**, **Figures S8**).

The free binding energy (∆G) of the RBD/hACE2 complex was calculated with Umbrella Sampling simulations. The profiles of the potential of mean force (PMF) along the reaction coordinate (ξ) at 150, 500, and 1,000 mM NaCl are shown in Figure 4. Free energy was calculated from the difference between the minimum and the maximum of the potential of mean force (PMF). Calculated dissociation constants (*K*_D_) were 0.7 and 1.0 nM at 150 and 1,000 mM NaCl, respectively (Table 1). However, this difference is not statistically significant. Consistent with the previous simulations, the Umbrella Sampling calculated *K*_D_ of 290 nM for RBD/hACE-2 complex at 500 mM NaCl is significantly higher than that determined at the other NaCl concentrations (Table 1).

**Table 1.** Free energy of Gibbs (*∆G*) and dissociation constant (*K*_D_) determined from Umbrella Sampling calculations performed in each salt concentration.

| NaCl concentration (mM) | $\Delta G$ (kcal.mol<sup>-1</sup>) | $K_D$ (nM) |
| --- | --- | --- |
| 150 | 13.0 ± 0.7 | 0.7 |
| 500 | 9.3 ± 0.8 | 290.0 |
| 1,000 | 12.8 ± 0.7 | 1.0 |

Equilibrium *K*_D_ were experimentally determined using surface plasmon resonance (SPR). Spike RBD/hACE-2 complex *K*_D_ were 1.1, 7.8 and 8.0 nM in the presence of 150, 500, and 1,000 mM NaCl, respectively (Table 2, **Table S9** and **Figure S9**), being in the same order of magnitude as the Umbrella Sampling data, particularly at 150 and 1,000 mM NaCl. Furthermore, complex affinity at 1,000 mM NaCl was slightly lower than that at 150 mM NaCl, which is consistent with the theoretical experiments. However, experimental *K*_D_ at 500 mM and 1,000 mM NaCl are nearly the same, which is in contrast to the computational experiments indicating a much lower binding affinity at this intermediate NaCl concentration. It is likely that intermolecular interactions become weaker at 500 mM, compared to 150 and 1,000 mM NaCl, which must be compensated by other effects, perhaps of entropic nature. However, these effects were not captured by simulations.

**Table 2.** Binding kinetics obtained by surface plasmon resonance experiments. Equilibrium dissociation constants (*K*_D_) were calculated for four spike constructs in the complex with hACE-2 in different NaCl concentrations.

| NaCl (mM) | RBD<sub>319-537</sub>/hACE-2<br>$K_D$ (nM) | *Trimer<sub>16-1213</sub>/hACE-2<br>$K_D$ (nM) | *Monomer<sub>16-1213</sub>/hACE-2<br>$K_D$ (nM) | S1<sub>16-685</sub>/hACE-2<br>$K_D$ (nM) |
| --- | --- | --- | --- | --- |
| 50 | 4.3 | 17.4 | n.d. | n.d. |
| 150 | 1.1 | 16.2 | n.d. | 12.3 |
| 500 | 7.8 | 8.0 | 17.8 | n.d. |
| 1,000 | 8.0 | 3.3 | 15.8 | 13.3 |

\*Monomeric and trimeric forms of Spike protein have substitutions at R683A and R685A; n.d.: not determined.

*K*_D_ of hACE-2 complexation with other Spike constructions were also measured experimentally (**Table S9** and **Figure S10-S12**). hACE-2 binds to the trimeric Spike with *K*_D_ = 16, 8, and 3 nM, in the presence of 150, 500, and 1,000 mM NaCl, respectively. In contrast to results obtained with RBD alone, the trimeric form Spike binds hACE-2 with higher affinity at 1,000 mM NaCl, when compared to 150 mM NaCl. High ionic strength supposedly favors the up orientation of the RBD within the trimeric form Spike. The other two Spike constructs, the S1 subunit, mimicking Spike fragment release during transition from prefusion to post-fusion states and the monomeric form of the full length Spike protein, showed only marginal differences in their *K*_D_ measured at different NaCl concentrations (Table 2). In all cases, association rate constants (*k*_a_ in **Table S9**, and *k* in **Table S6**) decrease with incrementing ionic strength, suggesting that long-range electrostatic interactions drive initial intermolecular attractions.

Overall, our experimental and theoretical data consistently demonstrate that a broad range of NaCl concentrations from 50 to 1,000 mM, does not disrupt the SARS-CoV-2 RBD/hACE-2 complex. At high ionic strengths, E2-mediated hydrophilic contacts become progressively weaker, while E1-mediated hydrophobic contacts increase to favor spike/hACE-2 complex maintenance. These features may reflect the necessity of complex stability during release of the S1 subunit that accompanies the structural transition of the Spike protein from the prefusion to the fusion intermediate state^18^.

## Supporting information

Dissociation_video-150mM

Dissociation_video-500mM

Dissociation_video-1000mM

