## supplemental information for "Molecular dynamics reveals complex compensatory effects of ionic strength on the SARS-CoV-2 Spike/hACE-2 interaction"

**Running title: Effects of NaCl concentration on the spike-ACE-2 interface**

To whom correspondence should be addressed:

Prof. Cristiane R. Guzzo, Department of Microbiology, Institute of Biomedical Sciences, University of São Paulo, Av. Prof. Lineu Prestes, 1374, Cidade Universitária, 5508-900, São Paulo/SP, Brazil, +55 11 3091-7298;

Prof. Roberto K. Salinas, Department of Biochemistry, Institute of Chemistry, University of São Paulo, Av. Prof. Lineu Prestes, 748, Cidade Universitária, 05508-000, São Paulo-SP, Brazil, +55 11 3091-1475;

**Keywords:** COVID-19, pandemic, coronavirus, spike, hACE-2.

### Materials and Methods

#### Molecular dynamics (MD) simulations

The initial coordinates for RBD/hACE-2 complex were obtained from the Protein Data Bank (PDB ID 6M0J, resolution 2.45 Å)[1] and prepared using the UCSF Chimera[2] by removing co-crystallized hetero groups (except $Zn^{2+}$) and water molecules. The system conditions were prepared using GROMACS v. 5.1.4[3–6] with the OPLS-AA force field[7]. We used the Swiss-Param web-based service to build the $Zn^{2+}$ topology[8]. All systems were explicitly solvated with *TIP3P* water models in a triclinic box (81.06 X 91.06 X 135.02 Å³), neutralized keeping NaCl concentration of 150 mM (114 $Na^+$ and 90 $Cl^-$ ions), 500 mM (324 $Na^+$ and 300 $Cl^-$ ions) and 1,000 mM (624 $Na^+$ and 600 $Cl^-$ ions) and minimized until a maximum force of 10.0 $kJ.mol^{-1}$ or 5,000 steps. The systems were consecutively equilibrated in isothermal-isochoric (NVT) and isothermal-isobaric (1 bar; NpT) ensembles at 310 K for 1000 ps. MD simulations in the NpT ensemble were carried out for 100 ns in a periodic cubic box considering the minimum distance of 10 Å between any protein atom and box walls. The $Zn^{2+}$ positions were restricted throughout the simulations. Distance calculations, root-mean-square deviation (RMSD), and root-mean-square fluctuation (RMSF) calculations were performed using GROMACS. The opening degree of the RBD/hACE-2 interface was quantified as the angle between two vectors chosen using the $C_\alpha$ coordinates along ACE-2 α-helix (residues 38 -49), and the Spike RBD β-strand (residues 492 - 494). Analysis of $C_\alpha$ cross-correlated displacements were performed using the R-based package Bio3d[9]. The covariances ranged from -1 (anti-correlated motions) to 1 (correlated motions). The $C_\alpha$ distances were calculated using GROMACS.

#### Hydrophobicity scale calculations

ClustalW/X[10] was used to compute multiple amino acid sequence alignments. The hydropathy indexes were calculated using Kyte and Doolitle[11].

#### Induced dissociation, free diffusion association and Umbrella Sampling simulation

The induced dissociation, free diffusion association, and Umbrella Sampling simulations were done with GROMACS 2018.6 [3–6] using the RBD/hACE-2 complex structure (PDB ID 6M0J, resolution 2.45 Å)[1] as initial coordinates and the CHARMM36 force field[12–14]. The systems were neutralized keeping the NaCl concentration at 150 mM (223 $Na^+$ and 199 $Cl^-$ ions), 500 mM (650 $Na^+$ and 626 $Cl^-$ ions) or 1,000 mM (1276 $Na^+$ and 1252 $Cl^-$ ions) in a triclinic box (100 X 100 X 210 $Å^3$) filled with 67,980, 62,954, and 61,752 *TIP3P* waters, respectively. The energy minimization was set to a maximum force of 1,000 kJ $mol^{-1}$ $nm^{-1}$ and a maximum number of steps of 50,000. The systems were equilibrated consecutively in isothermal-isochoric (NVT) and isothermal-isobaric (1 bar; NpT) ensembles at 310 K for 2000 ps, followed by the calculation of a 100 ns MD trajectory in the NpT ensemble.

*Induced dissociation*

Starting from the last frame of 100 ns MD trajectory, a reaction coordinate was calculated as the average of the $C_\alpha$ (RBD) - $C_\alpha$ (hACE-2) intermolecular vectors of all residue pairs at the interface[1]. The intermolecular vectors were computed using an in-house python script. The RBD/hACE-2 complex was rotated to align the z-axis of the structure with the reaction coordinate vector. A harmonic potential of 600 kJ $mol^{-1}.nm^{-2}$ constant force and 0.005 $nm.ns^{-1}$ pulling speed was applied on the center of mass of the RBD, while the positions of the hACE-2 atoms were restricted using a harmonic force of 1000 $kJ.mol^{-1}.nm^{-2}$. The pulling trajectory time was 1 ns.

*Free diffusion simulation*

The RBD and hACE-2 were pulled as described above up to a center-of-mass distance of ~60 Å. The separation distance should be large to capture diffusion effects, but not as large to avoid interactions between them. To equilibrate the system, it was relaxed during a 3 ns annealing, in which the temperature was increased from 310 K to 350 K in 500 ps, followed by an NpT equilibrium dynamics at 350 K for 2 ns, and a temperature decrease from 350 K to 310 K in 500 ps. The two molecules were then released of all restriction forces, and a MD simulation of 20 ns was performed. When this experiment was performed at 500 mM NaCl, there was no recovery of the RBD/hACE-2 complex after 20 ns of simulation. The analysis was based on the 2,000 snapshots saved every 10 ps during the trajectory. The number of $Na^+$ and $Cl^-$ ions within a radius of 6 Å from each amino acid at the binding interface was computed using an in-house python script.

*Umbrella Sampling*

Initial configurations for a biased sampling simulation were recorded at every step of ~ 0.1 nm in the dissociation pathway. Umbrella sampling (US) windows were equilibrated in the NpT ensemble using the simulated annealing protocol described above, followed by 10 ns simulation. In total, 33 configurations were calculated at 150 and 500 mM NaCl concentrations, while 30 configurations were computed at 1,000 mM NaCl solution. The three additional configurations at 150 and 500 mM NaCl concentrations generated overlapping biased distribution functions. The weighted histogram analysis method (WHAM)[15] was used to compute the potential of mean force (PMF) along the reaction coordinate (ξ) with the g_wham GROMACS module. Free binding energies (ΔG) were estimated from the difference between the minimum and maximum of the PMF profile in a given NaCl concentration. Dissociation constants ($K_D$) were determined assuming $\Delta G = RTln\,(K_D)$, $R$ and $T$ are gas constant and temperature, respectively. PMF errors were calculated using 100 rounds of bootstrapping analysis.

**Protein-protein binding assays using surface plasmon resonance (SPR)**

The experimental $K_D$ ($M^{-1}$) of the spike/hACE-2 complex were obtained by surface plasmon resonance (SPR) assays using a Biacore 8K system (GE Healthcare). The assays as well as protein cloning, expression and purification were performed by the ACROBiosystems company. In all SPR experiments were used four different spike protein constructs: the RBD (residues 319 to 537); the S1 subunit (residues 16 to 685); monomeric form of the spike protein (residues 16 to 1213, with mutations at arginines 683 and 685 to alanines: R683A, R685A); and trimeric form of spike protein (residues

16 to 1213, with mutations at arginines 683 and 685 to alanines: R683A, R685A). Two different recombinant hACE-2 (residues from 18 - 740) constructs were used: hACE-2 with C-terminal Fc tag (MW 110 kDa; AC2-H5257, ACROBiosystems) and biotinylated hACE-2 with C-terminal polyhistidine and Avi tags (MW 87.2 kDa; AC2-H82E6, ACROBiosystems), in which both hACE-2 were dimers. Anti-Human IgG (Fc) antibody was diluted at 25 µg/mL in 10 mM Sodium Acetate, pH 5.0, to be covalently immobilized to a CM5 sensor chip (GE Healthcare) via their amine groups using the human antibody capture kit (BR-1008-39, GE Healthcare). Immobilization processes were performed using a flow rate of Anti-Human IgG (Fc) antibody of 10 $\mu L.min^{-1}$ during 420 s, where obtained response unit (RU) signals were about 9,000-14,000. Human ACE-2 with a C-terminal Fc tag was diluted with running buffers (10 mM HEPES, pH 7.5, 0.05% Tween, supplemented with 50 - 1,000 mM NaCl). The RU signal obtained was about 150 in the capture hACE-2 protein-ligand. After that, serial dilutions of purified recombinant Spike proteins were injected ranging in concentrations from 2 to 15 nM for SARS-CoV-2 RBD (MW 26.5 kDa; SPD-C52H3, ACROBiosystems), from 4 to 250 nM for S1 subunit (MW 76.9 kDa; S1N-C52H2, ACROBiosystems), and from 2 to 500 nM for a trimeric form of Spike (MW 138.1 kDa; SPN-C52H8, ACROBiosystems). Spike constructs were injected at a flow rate of 30 $\mu L.min^{-1}$ during association (120 s) and dissociation (200 s). The chip was regenerated using a regeneration solution composed of a mix of 3 parts of 8 M guanidine hydrochloride and 1 part of 1 M sodium hydroxide (GE) at a flow rate of 20 $\mu L.min^{-1}$ in 30 s. The resulting data were fitted to a 1:1 binding model using Biacore Evaluation Software (GE Healthcare). A similar assay was performed to calculate $K_D$ values for complexes of biotinylated hACE-2 and monomeric form of the Spike. Initially, Series S Sensor Chip CAP (GE Healthcare) were prepared using the Biotin capture kit (Series S, GE Healthcare) to capture biotinylated molecules. The immobilization process was performed using a flow rate of 2 $\mu L.min^{-1}$ during 60 s, where RU signal levels were about 3,000. Biotinylated hACE-2 at the final concentration of 10 $\mu g.mL^{-1}$ was diluted in different running buffers as described above. During the ligand capture process, a flow rate of 10 $\mu L.min^{-1}$ was used until the RU signal reached a level of ~100. After that, serial dilutions of purified recombinant Spike were injected ranging in concentration from 4 to 25 nM for monomeric form of Spike (MW 134.6 kDa; SPN-C52H4, ACROBiosystems). The Spike constructs were injected at a flow rate of 30

μL.min$^{-1}$ during association (120s) and dissociation (200s) processes. The chip was regenerated using a regeneration solution composed of a mix of 3 parts of 8M guanidine hydrochloride and 1 part of 1 M sodium hydroxide (GE) at a flow rate of 10 μL.min$^{-1}$ during 120 s. Resulting data were fitted to a 1:1 binding model using Biacore Evaluation Software (GE Healthcare).

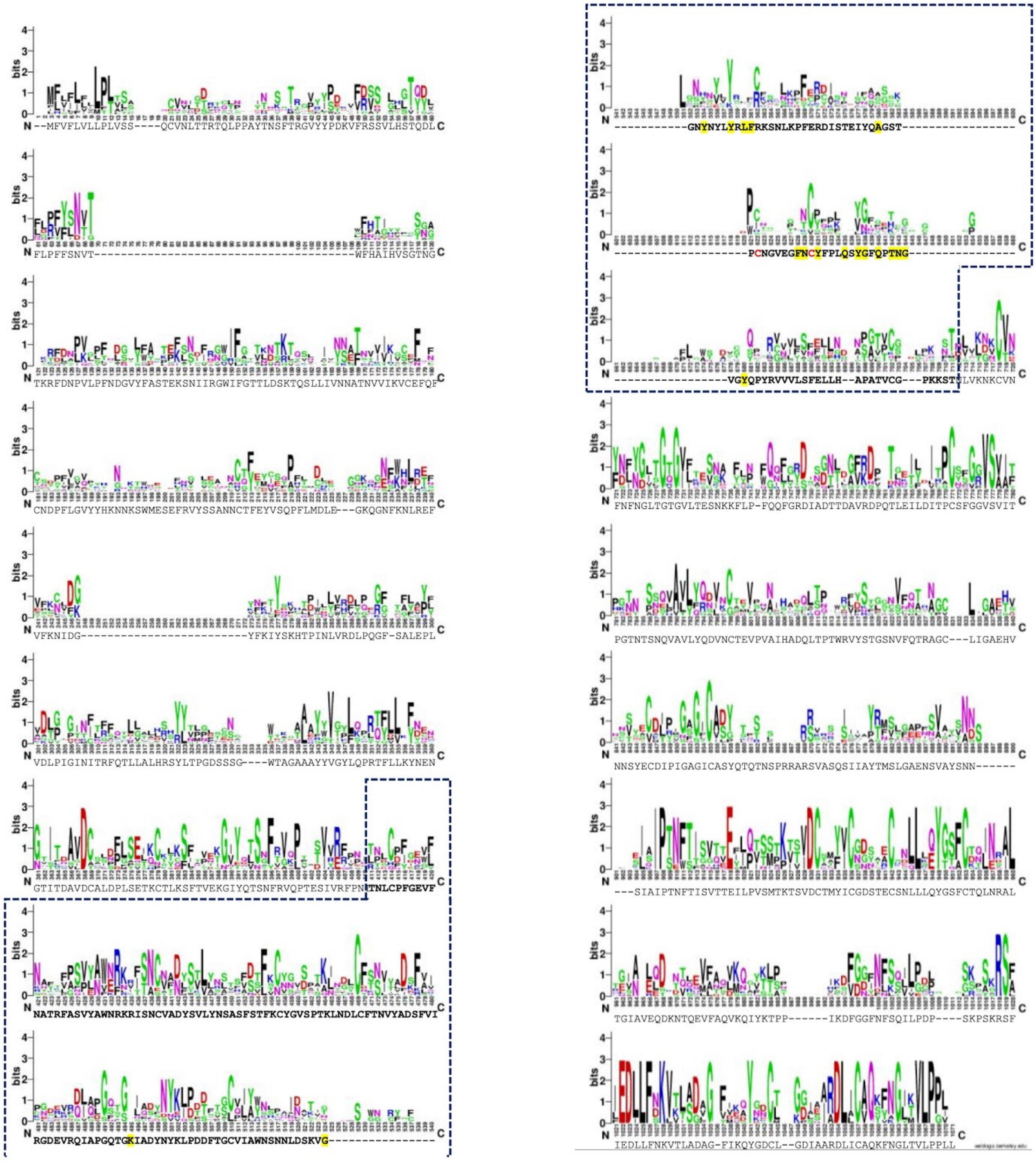

**Figure S1:** Multiple amino acid sequence alignment of Spike S1 subunit within coronavirus members is represented by WebLogo. The sequence shown below the logo belongs to SARS-CoV-2. The residues in bold belong to RBD and residues highlighted in yellow interact with the human ACE-2 receptor[1]. The cysteines colored in red form a thiosulfate bond in the E1 region probably to maintain the $\beta$7-$\beta$8 hairpin stable in the three-dimensional structure (shown in the **Figure 1c** in the main text).

The Spike sequences used are of SARS-CoV (GenBank: AAP41037.1), SARS-CoV-2 (from a patient from Brazil, GenBank: QIG55994.1), others human coronavirus (from a patient from United States in 2017, GenBank: AZS52618.1, and from Hong Kong in 2006, GenBank: ABD75529.1), HCoV229E (Gene ID: 918758), HcoV_NL63 (Gene ID:2943499), HcoV-OC43 (Gene ID: 39105218), and Middle East respiratory syndrome-related coronavirus, MERS (Gene ID YP_009047204.1). We also used the sequence of bat RaTG13 (GenBank: QHR63300.2) and pangolin coronaviruses (GenBank: QIA48632.1). The multiple sequence alignment was created using ClustalW[16]. The sequence logo was created with the WebLogo server[17]. The residues belonging to the RBD are inside of the dotted blue square.

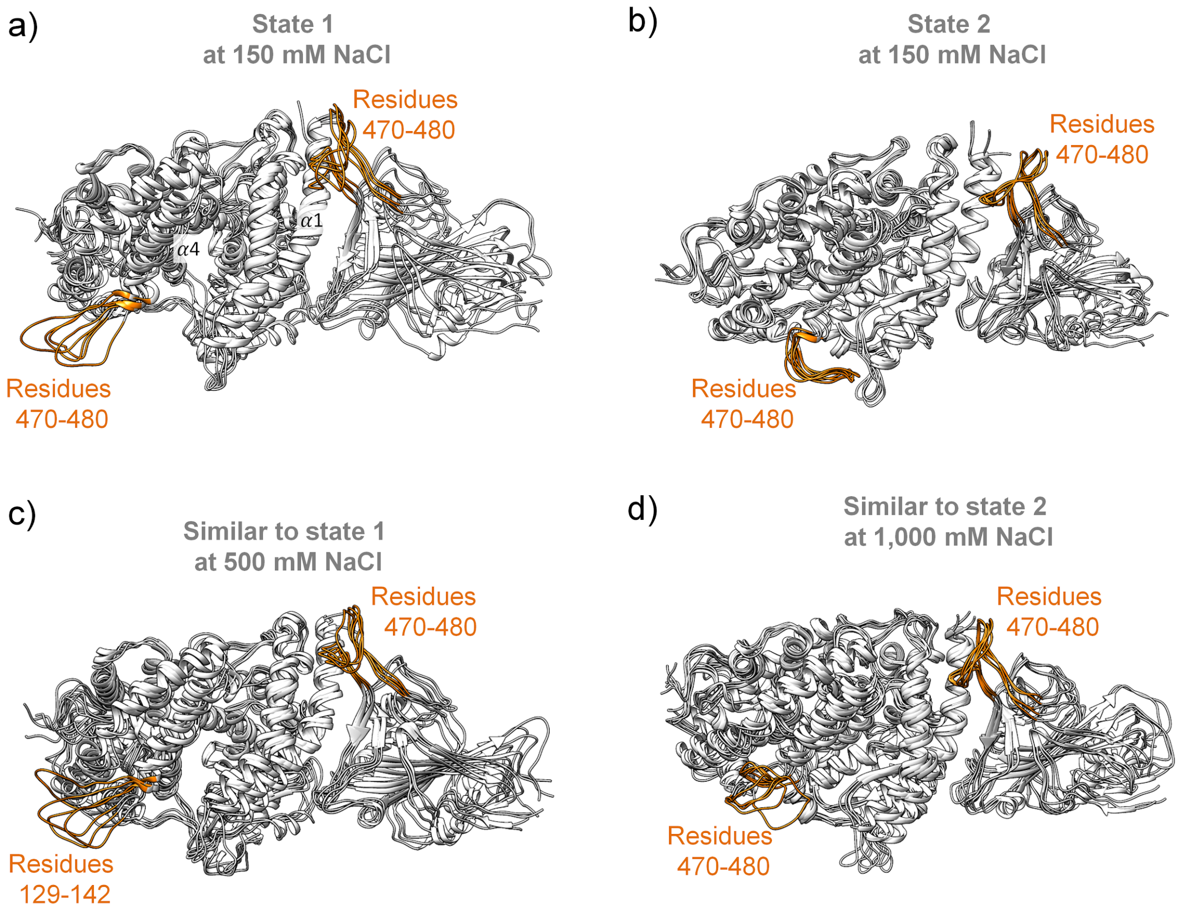


**Figure S2.** Conformational states associated with RBD loop 470-490 and hACE-2 loop 129-142 (both highlighted in yellow). RBD/hACE-2 visited two conformational states at 150 mM NaCl: (a) state 1 and (b) state 2. At 500 mM NaCl, the complex assumes a conformation close to state 1 (c). In the presence of 1,000 mM NaCl, the complex conformation is similar to state 2 (d).

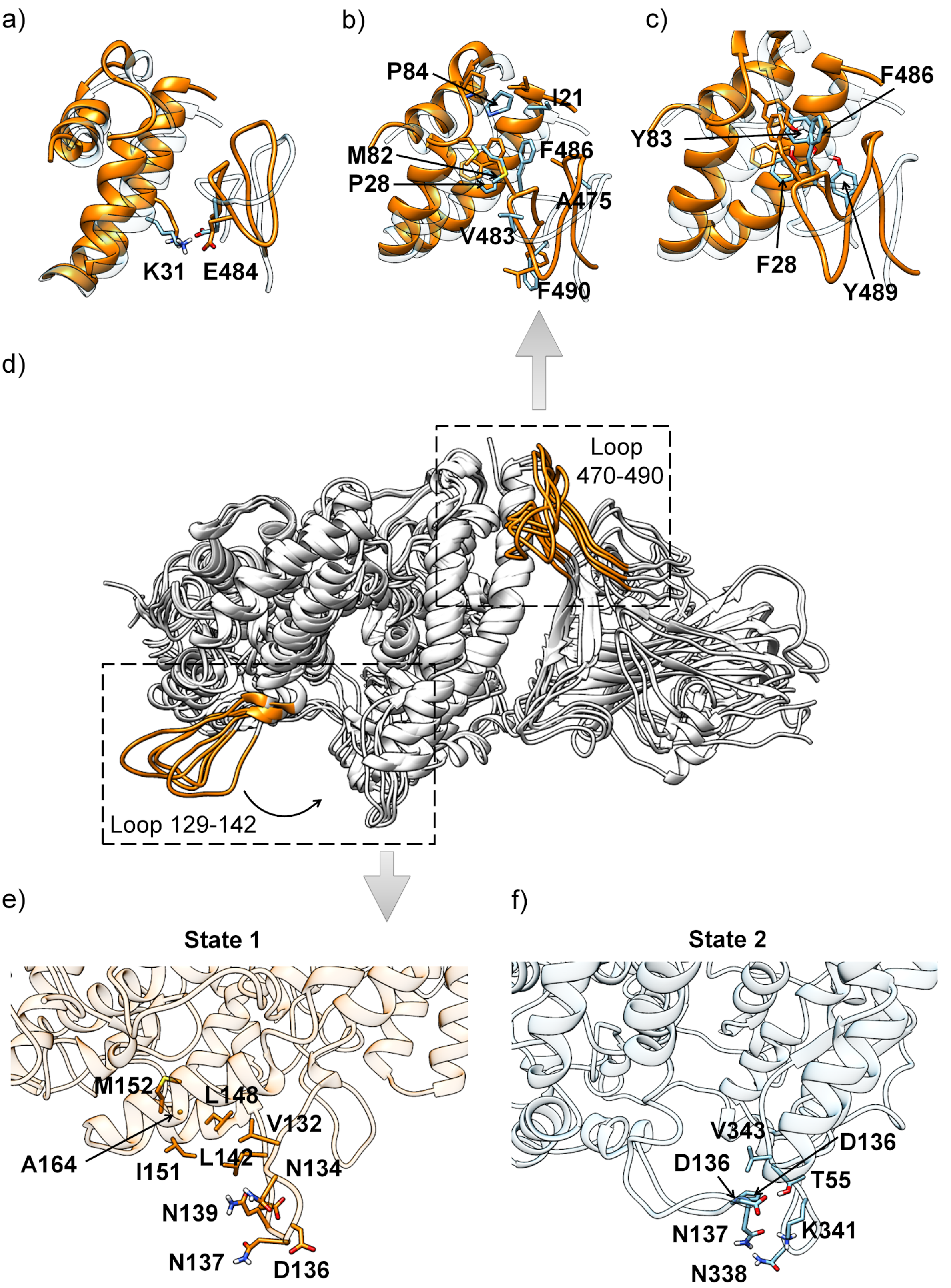


**Figure S3.** E1 region/hACE-2 interactions. (a) π-stacking interactions between residue pairs $F486^{RBD}/Y83^{hACE-2}$ and $Y489^{RBD}/F28^{hACE-2}$, (b) van der Waals interactions and (c) salt bridge between $K31^{hACE-2}$ and $E484^{RBD}$. (d) hACE-2 may visit states 1 and 2. The transition from state 1 to 2 is characterized by $\sim 90^0$ rotation of residues 129-142 to interact with $\alpha 1$ - $\alpha 2$ (residues 50-55) and $\alpha 10$ - $\beta 4$ (residues 330-347) loops. (e) State 1 is characterized by hydrogen bonding interactions between residue pairs

N134/N137, N137/D136, Q139/E140 and the juxtaposition of the hydrophobic V132/L142 in the vicinity of the hydrophobic region formed by I151, M152, L148, and A164 residues. (f) State 2 is stabilized by hydrogen bonding pairs D136/T55 and N137/N338, nonpolar contacts between P138 and V343, and a salt bridge between D136 and K341.

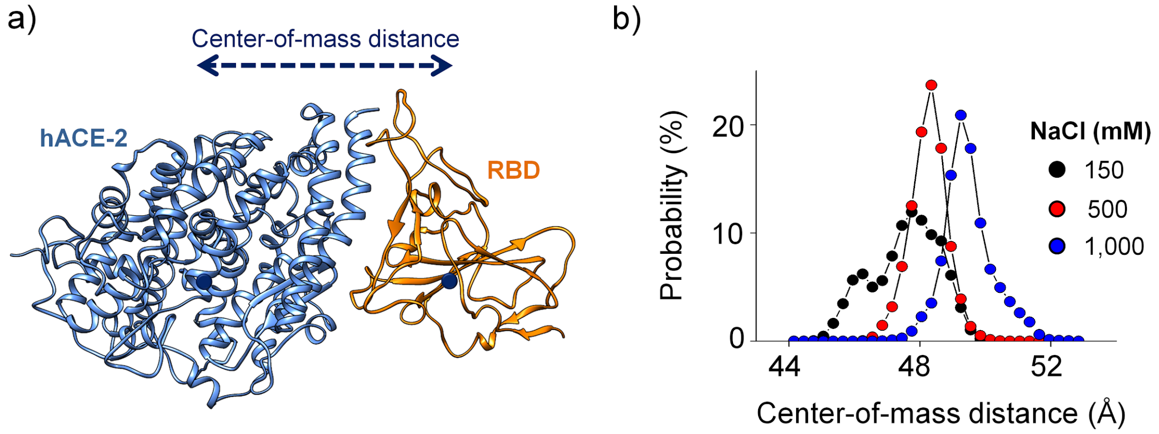


**Figure S4.** (a) Spacing of the interaction interface as a function of the increment in NaCl concentration. (b) Probability distribution of center-of-mass distance at 150 mM (black), 500 mM (red), and 1,000 mM (blue) NaCl concentrations.

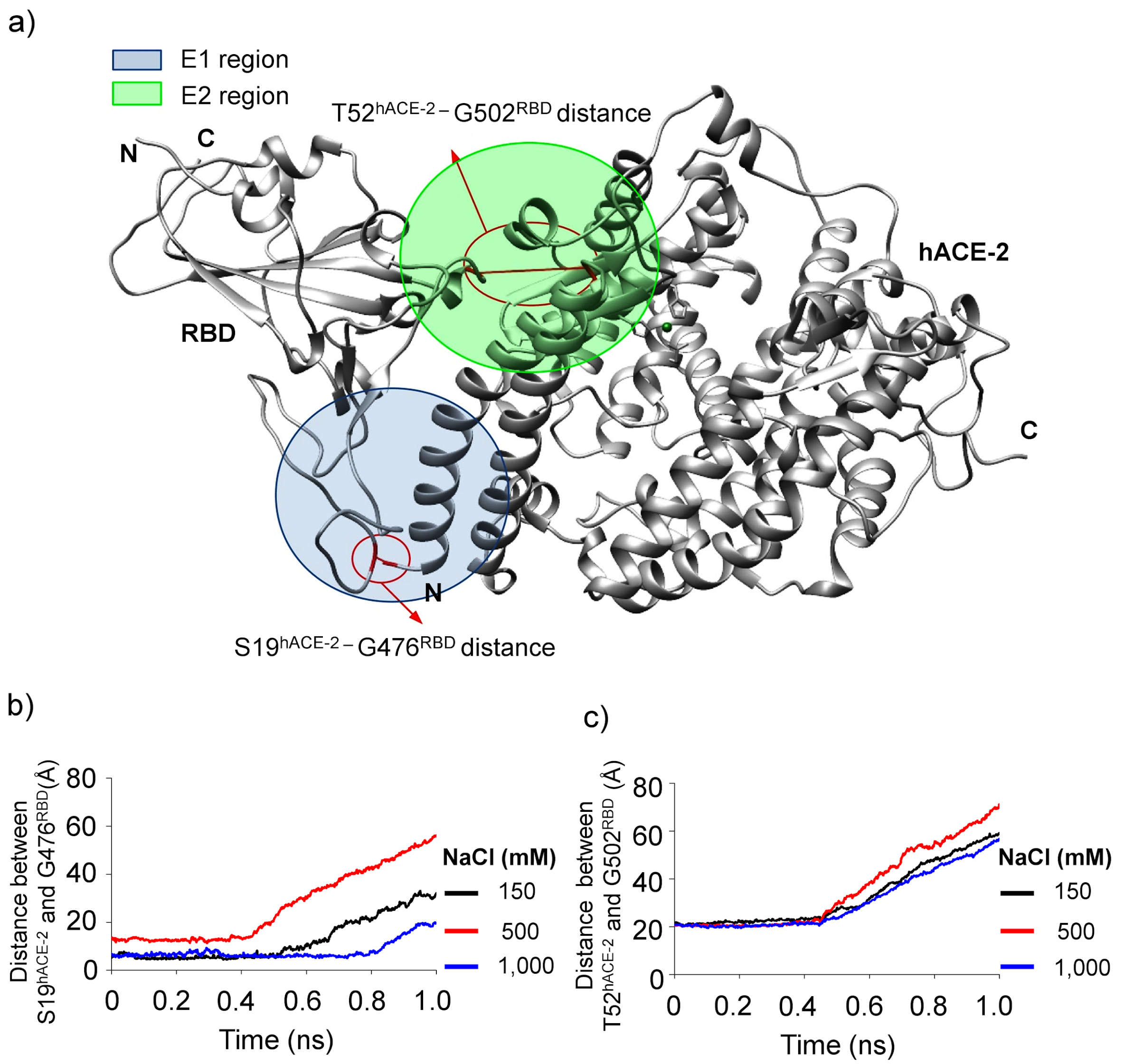


**Figure S5.** Induced dissociation of the RBD/hACE-2 complex. **a**) The two distances that were monitored during the simulations are indicated: G476$^{RBD}$-S19$^{hACE\text{-}2}$ in E1 and G502$^{RBD}$-T52$^{hACE\text{-}2}$ in E2. **b)** The initial S19$^{hACE\text{-}2}$-G476$^{RBD}$ distance of ~5-6 Å displays a longer lag phase at 1,000 mM when compared to 150 mM NaCl, a difference of ~0.3 ns. At 500 mM, the initial distance is ~13 Å, and the lag phase is significantly shorter, ~0.4 ns. **b**) The T52$^{hACE\text{-}2}$-G502$^{RBD}$ distances are maintained in all salt concentrations until ~0.45 ns, after what the separation begins. The separation rate is slighter faster at 500 mM when compared to the other NaCl concentrations.

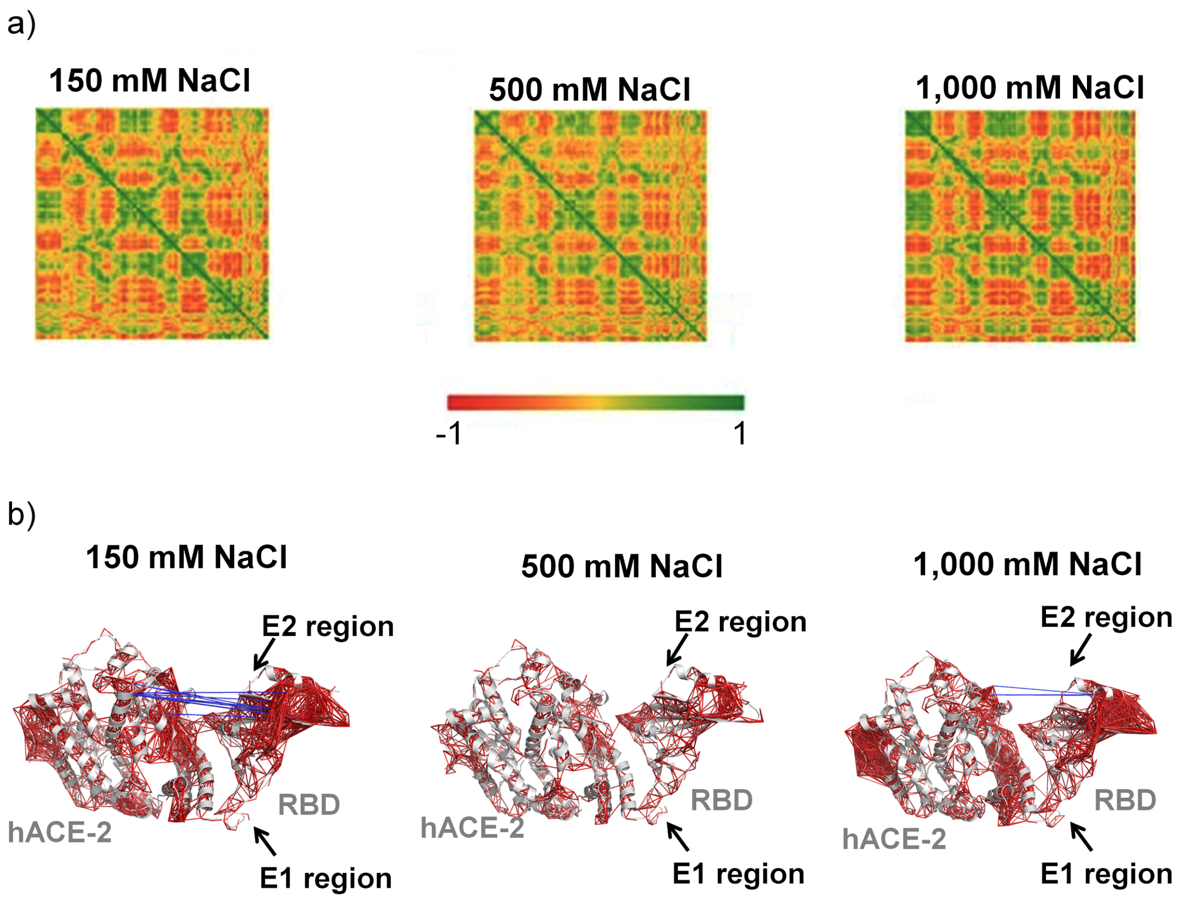


**Figure S6.** $C_\alpha$ spatial displacement covariances during the MD analysis of RBD/hACE-2 in increasing salt concentrations. **a**) Covariance matrices between $C_\alpha$ versus $C_\alpha$ considering hACE-2 and RBD sequences. **b**) Pairs of $C_\alpha$ exhibiting covariances greater than 0.8 (positive and negative) were mapped in the Spike RBD/hACE-2 complex three-dimensional structure (PDB ID 6M0J[1]). Red lines mark positive covariance pairs, whereas blue lines show negative ones.

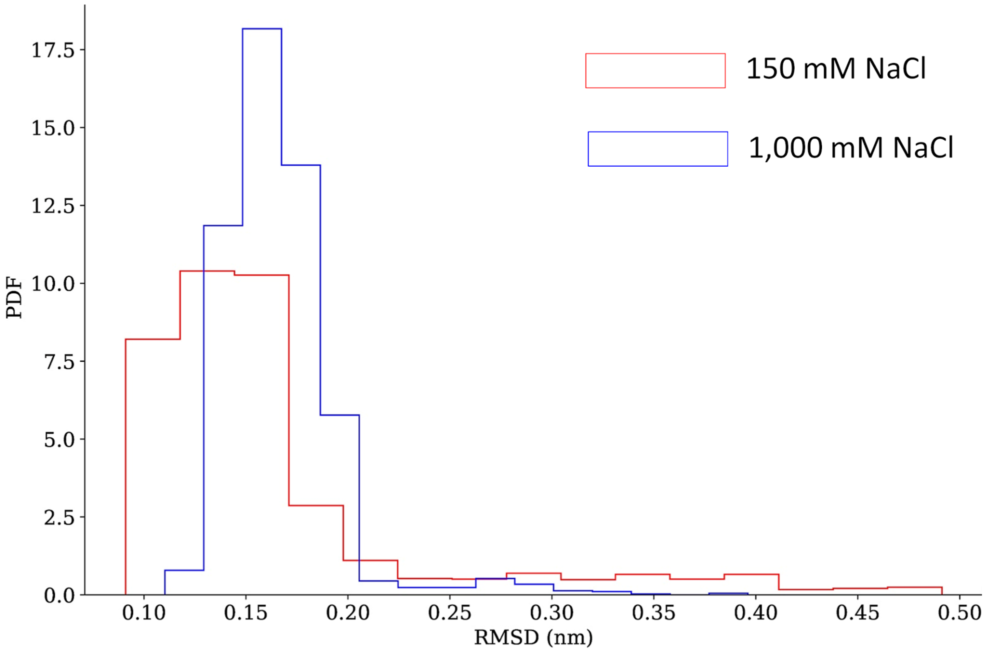


**Figure S7.** Probability distribution functions (PDF) of all atom-RMSD at the RBD/hACE-2 interface in the free diffusion trajectories. The representative RMSD at 150 and 1,000 mM NaCl is 1 - 2 Å, indicating the association of the RBD/hACE-2 complex.

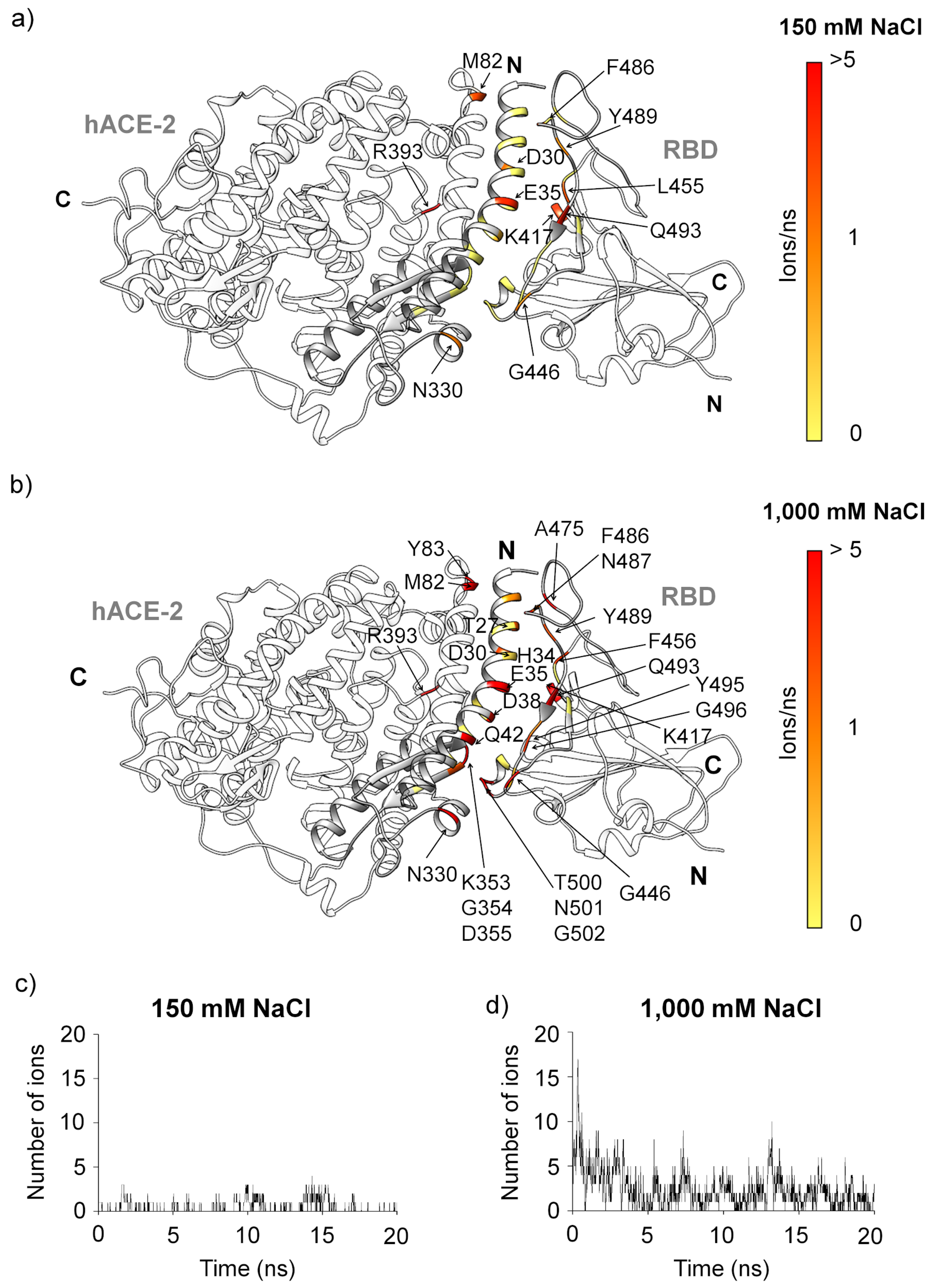


**Figure S8**. Number of ions per nanosecond within 6 Å from each amino acid along the diffusion trajectory color coded on the crystal structure of SARS-CoV-2 RBD/h-ACE2 (PDB ID: 6M0J[1]). The amino acids displaying more than one ion per nanosecond in their vicinity are indicated: **(a)** 150 mM NaCl and **(b)** 1,000 mM NaCl; Sum of the number of ions at the interface during the trajectory at **(c)** 150 mM NaCl and **(d)** 1,000 mM NaCl.

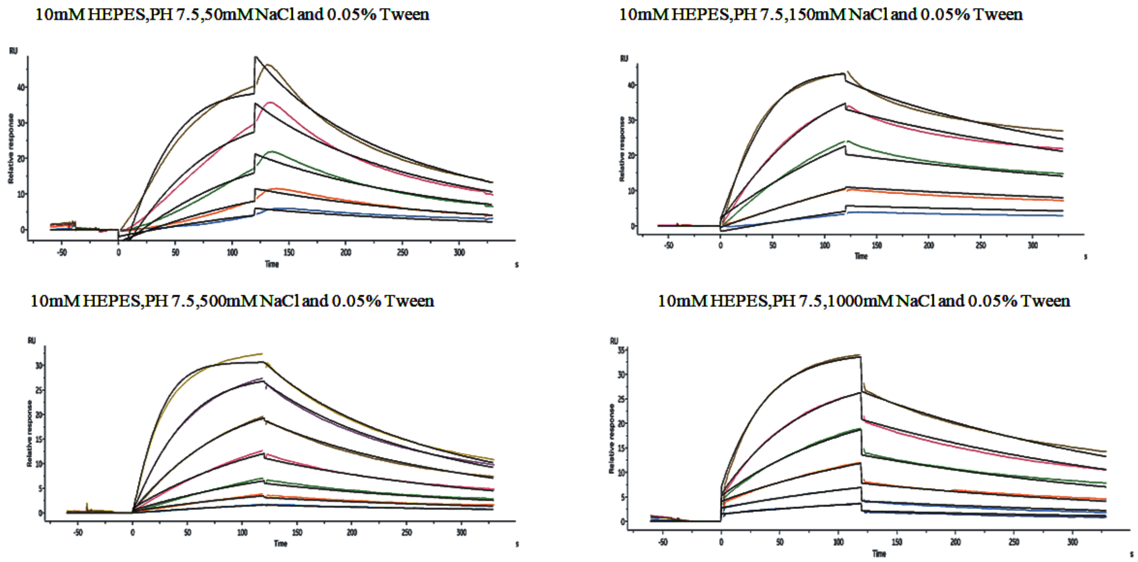


**Figure S9.** Surface plasmon resonance (SPR) data using immobilized recombinant dimer hACE-2 and different concentrations of purified recombinant Spike RBD (residues residues 319 to 537), ranging from 2 to 15 nM. The experimental data were fitted to a 1:1 binding model using Biacore Evaluation Software (GE Healthcare) to calculate the kinetics parameters ($k_a$ and $k_d$) shown in **Table 2** in the main text and **Table S9**. At least five different RBD concentrations were used to calculate the $K_D$ values.

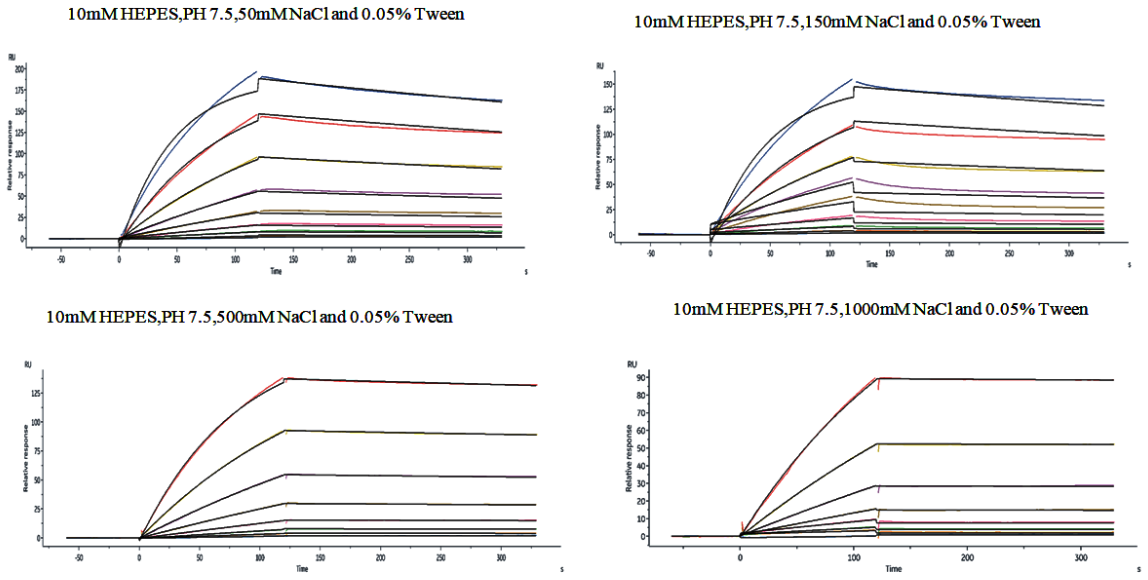

**Figure S10.** Surface plasmon resonance (SPR) data using immobilized recombinant dimer hACE-2 and different concentrations of purified recombinant trimeric form of the Spike (residues 16 to 1213, with mutations at arginines 683 and 685 to alanines: R683A, R685A), ranging from 2 to 500 nM. The experimental data were fitted to a 1:1 binding model using Biacore Evaluation Software (GE Healthcare) to calculate the kinetics parameters ($k_a$ and $k_d$) shown in **Table 2** in the main text and **Table S9**. At least six different trimeric form of the Spike concentrations were used to calculate the $K_D$ values.

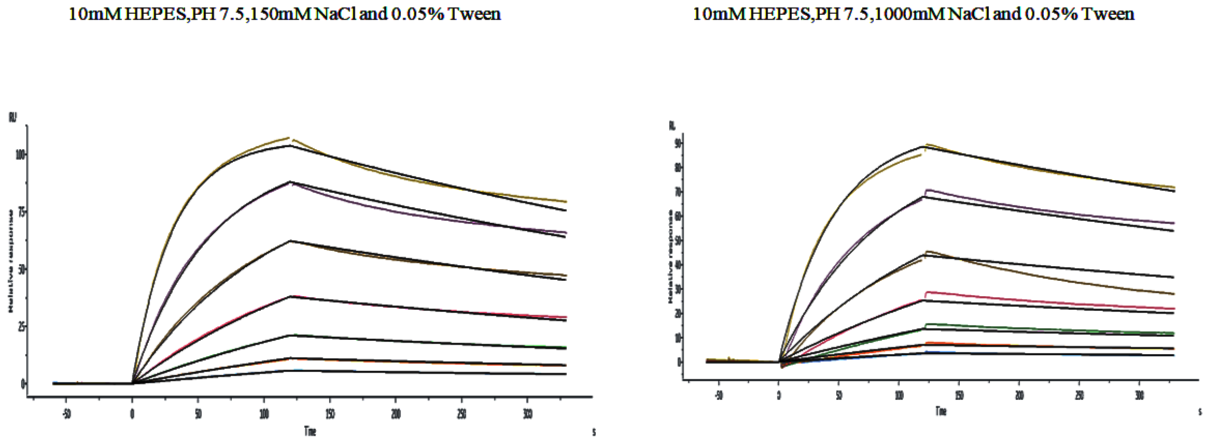


**Figure S11.** Surface plasmon resonance (SPR) data using immobilized recombinant dimer hACE-2 and different concentrations of purified recombinant S1 subunit of Spike (residues residues 16 to 685), ranging from 4 to 250 nM. The experimental data were fitted to a 1:1 binding model using Biacore Evaluation Software (GE Healthcare) to calculate the kinetics parameters ($k_a$ and $k_d$) shown in **Table 2** in the main text and **Table S9**. At least five different spike S1 subunit concentrations were used to calculate the $K_D$ values.

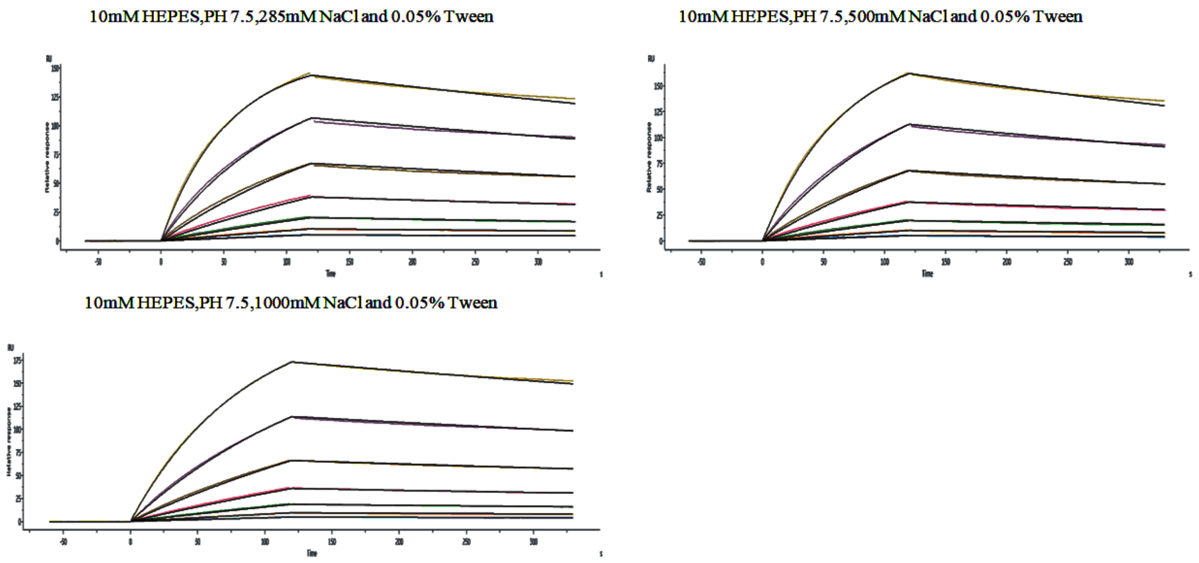

**Figure S12.** Surface plasmon resonance (SPR) data using immobilized recombinant dimer hACE-2 and different concentrations of purified recombinant monomeric form of the Spike (residues 16 to 1213, with mutations at arginines 683 and 685 to alanines: R683A, R685A), ranging from 4 to 25 nM. The experimental data were fitted to a 1:1 binding model using Biacore Evaluation Software (GE Healthcare) to calculate the kinetics parameters ($k_a$ and $k_d$) shown in **Table 2** in the main text and **Table S9**. Seven different Spike concentrations were used to calculate the $K_D$ values.

**Table S1.** The hydropathy scale calculated from Kyte and Doolittle (1982)[11] for each residue of RBDs of SARS-CoV and SARS-CoV-2. Residues located in the E1 region are colored in green and those located in the E2 region in orange. Residues in asterisks make part of the hACE-2 interface.

| Amino acid residues | | | Normalized hydropathy | | |
|---|---|---|---|---|---|
| number | SARS-CoV | SARS-CoV 2 | SARS-CoV | SARS-CoV 2 | Δ hydropathy (SARS-CoV-2 take way SARS-CoV) |
| 333 | T | T | -0.7 | -0.7 | 0 |
| 334 | N | N | -3.5 | -3.5 | 0 |
| 335 | L | L | 3.8 | 3.8 | 0 |
| 336 | C | C | 2.5 | 2.5 | 0 |

| | | | | | |
|---|---|---|---|---|---|
| 337 | P | P | -1.6 | -1.6 | 0 |
| 338 | F | F | 2.8 | 2.8 | 0 |
| 339 | G | G | -0.4 | -0.4 | 0 |
| 340 | E | E | -3.5 | -3.5 | 0 |
| 341 | V | V | 4.2 | 4.2 | 0 |
| 342 | F | F | 2.8 | 2.8 | 0 |
| 343 | N | N | -3.5 | -3.5 | 0 |
| 344 | A | A | 1.8 | 1.8 | 0 |
| 345 | T | T | -0.7 | -0.7 | 0 |
| 346 | K | R | -3.9 | -4.5 | -0.6 |
| 347 | F | F | 2.8 | 2.8 | 0 |
| 348 | P | A | -1.6 | 1.8 | 3.4 |
| 349 | S | S | -0.8 | -0.8 | 0 |
| 350 | V | V | 4.2 | 4.2 | 0 |
| 351 | Y | Y | -1.3 | -1.3 | 0 |
| 352 | A | A | 1.8 | 1.8 | 0 |
| 353 | W | W | -0.9 | -0.9 | 0 |
| 354 | E | N | -3.5 | -3.5 | 0 |
| 355 | R | R | -4.5 | -4.5 | 0 |

| | | | | | |
|---|---|---|---|---|---|
| 356 | K | K | -3.9 | -3.9 | 0 |
| 357 | K | R | -3.9 | -4.5 | -0.6 |
| 358 | I | I | 4.5 | 4.5 | 0 |
| 359 | S | S | -0.8 | -0.8 | 0 |
| 360 | N | N | -3.5 | -3.5 | 0 |
| 361 | C | C | 2.5 | 2.5 | 0 |
| 362 | V | V | 4.2 | 4.2 | 0 |
| 363 | A | A | 1.8 | 1.8 | 0 |
| 364 | D | D | -3.5 | -3.5 | 0 |
| 365 | Y | Y | -1.3 | -1.3 | 0 |
| 366 | S | S | -0.8 | -0.8 | 0 |
| 367 | V | V | 4.2 | 4.2 | 0 |
| 368 | L | L | 3.8 | 3.8 | 0 |
| 369 | Y | Y | -1.3 | -1.3 | 0 |
| 370 | N | N | -3.5 | -3.5 | 0 |
| 371 | S | S | -0.8 | -0.8 | 0 |
| 372 | T | A | 0.7 | 1.8 | 1.1 |
| 373 | F | S | 2.8 | -0.8 | -3.6 |
| 374 | F | F | 2.8 | 2.8 | 0 |

| | | | | | |
|---|---|---|---|---|---|
| 375 | S | S | -0.8 | -0.8 | 0 |
| 376 | T | T | -0.7 | -0.7 | 0 |
| 377 | F | F | 2.8 | 2.8 | 0 |
| 378 | K | K | -3.9 | -3.9 | 0 |
| 379 | C | C | 2.5 | 2.5 | 0 |
| 380 | Y | Y | -1.3 | -1.3 | 0 |
| 381 | G | G | -0.4 | -0.4 | 0 |
| 382 | V | V | 4.2 | 4.2 | 0 |
| 383 | S | S | -0.8 | -0.8 | 0 |
| 384 | A | P | 1.8 | -1.6 | -3.4 |
| 385 | T | T | -0.7 | -0.7 | 0 |
| 386 | K | K | -3.9 | -3.9 | 0 |
| 387 | L | L | 3.8 | 3.8 | 0 |
| 388 | N | N | -3.5 | -3.5 | 0 |
| 389 | D | D | -3.5 | -3.5 | 0 |
| 390 | L | L | 3.8 | 3.8 | 0 |
| 391 | C | C | 2.5 | 2.5 | 0 |
| 392 | F | F | 2.8 | 2.8 | 0 |

| | | | | | |
|---|---|---|---|---|---|
| 393 | S | T | -0.8 | 0.7 | 1.5 |
| 394 | N | N | -3.5 | -3.5 | 0 |
| 395 | V | V | 4.2 | 4.2 | 0 |
| 396 | Y | Y | -1.3 | -1.3 | 0 |
| 397 | A | A | 1.8 | 1.8 | 0 |
| 398 | D | D | -3.5 | -3.5 | 0 |
| 399 | S | S | -0.8 | -0.8 | 0 |
| 400 | F | F | 2.8 | 2.8 | 0 |
| 401 | V | V | 4.2 | 4.2 | 0 |
| 402 | V | I | 4.2 | 4.5 | 0.3 |
| 403 | K | R | -3.9 | -4.5 | -0.6 |
| 404 | G | G | -0.4 | -0.4 | 0 |
| 405 | D | D | -3.5 | -3.5 | 0 |
| 406 | D | E | -3.5 | -3.5 | 0 |
| 407 | V | V | 4.2 | 4.2 | 0 |
| 408 | R | R | -4.5 | -4.5 | 0 |
| 409 | Q | Q | -3.5 | -3.5 | 0 |
| 410 | I | I | 4.5 | 4.5 | 0 |
| 411 | A | A | 1.8 | 1.8 | 0 |

| | | | | | |
|---|---|---|---|---|---|
| 412 | P | P | -1.6 | -1.6 | 0 |
| 413 | G | G | -0.4 | -0.4 | 0 |
| 414 | Q | Q | -3.5 | -3.5 | 0 |
| 415 | T | T | -0.7 | -0.7 | 0 |
| 416 | G | G | -0.4 | -0.4 | 0 |
| 417 | V | K* | 4.2 | -3.9 | -8.1 |
| 418 | I | I | 4.5 | 4.5 | 0 |
| 419 | A | A | 1.8 | 1.8 | 0 |
| 420 | D | D | -3.5 | -3.5 | 0 |
| 421 | Y | Y | -1.3 | -1.3 | 0 |
| 422 | N | N | -3.5 | -3.5 | 0 |
| 423 | Y | Y | -1.3 | -1.3 | 0 |
| 424 | K | K | -3.9 | -3.9 | 0 |
| 425 | L | L | 3.8 | 3.8 | 0 |
| 426 | P | P | -1.6 | -1.6 | 0 |
| 427 | D | D | -3.5 | -3.5 | 0 |
| 428 | D | D | -3.5 | -3.5 | 0 |
| 429 | F | F | 2.8 | 2.8 | 0 |

| | | | | | |
|---|---|---|---|---|---|
| 430 | M | T | 1.9 | 0.7 | -1.2 |
| 431 | G | G | -0.4 | -0.4 | 0 |
| 432 | C | C | 2.5 | 2.5 | 0 |
| 433 | V | V | 4.2 | 4.2 | 0 |
| 434 | L | I | 3.8 | 4.5 | 0.7 |
| 435 | A | A | 1.8 | 1.8 | 0 |
| 436 | W | W | -0.9 | -0.9 | 0 |
| 437 | N | N | -3.5 | -3.5 | 0 |
| 438 | T | S | 0.7 | -0.8 | -1.5 |
| 439 | R | N | -4.5 | -3.5 | 1 |
| 440 | N | N | -3.5 | -3.5 | 0 |
| 441 | I | L | 4.5 | 3.8 | -0.7 |
| 442 | D | D | -3.5 | -3.5 | 0 |
| 443 | A | S | 1.8 | -0.8 | -2.6 |
| 444 | T | K | 0.7 | -3.9 | -4.6 |
| 445 | S | V | -0.8 | 4.2 | 5 |
| 446 | T | G* | 0.7 | -0.4 | -1.1 |
| 447 | G | G | -0.4 | -0.4 | 0 |
| 448 | N | N | -3.5 | -3.5 | 0 |

| | | | | | |
|---|---|---|---|---|---|
| 449 | Y | Y* | -1.3 | -1.3 | 0 |
| 450 | N | N | -3.5 | -3.5 | 0 |
| 451 | Y | Y | -1.3 | -1.3 | 0 |
| 452 | K | L | -3.9 | 3.8 | 7.7 |
| 453 | Y | Y* | -1.3 | -1.3 | 0 |
| 454 | R | R | -4.5 | -4.5 | 0 |
| 455 | Y | L* | -1.3 | 3.8 | 5.1 |
| 456 | L | F* | 3.8 | 2.8 | -1 |
| 457 | R | R | -4.5 | -4.5 | 0 |
| 458 | H | K | -3.2 | -3.9 | -0.7 |
| 459 | G | S | -0.4 | -0.8 | -0.4 |
| 460 | K | N | -3.9 | -3.5 | 0.4 |
| 461 | L | L | 3.8 | 3.8 | 0 |
| 462 | R | K | -4.5 | -3.9 | 0.6 |
| 463 | P | P | -1.6 | -1.6 | 0 |
| 464 | F | F | 2.8 | 2.8 | 0 |
| 465 | E | E | -3.5 | -3.5 | 0 |
| 466 | R | R | -4.5 | -4.5 | 0 |
| 467 | D | D | -3.5 | -3.5 | 0 |

| | | | | | |
|---|---|---|---|---|---|
| 468 | I | I | 4.5 | 4.5 | 0 |
| 469 | S | S | -0.8 | -0.8 | 0 |
| 470 | N | T | -3.5 | 0.7 | 4.2 |
| 471 | V | E | 4.2 | -3.5 | -7.7 |
| 472 | P | I | -1.6 | 4.5 | 6.1 |
| 473 | F | Y | 2.8 | -1.3 | -4.1 |
| 474 | S | Q | -0.8 | -3.5 | -2.7 |
| 475 | P | A* | -1.6 | 1.8 | 3.4 |
| 476 | D | G | -3.5 | -0.4 | 3.1 |
| 477 | G | S | -0.4 | -0.8 | -0.4 |
| 478 | K | T | -3.9 | 0.7 | 4.6 |
| 479 | P | P | -1.6 | -1.6 | 0 |
| 480 | C | C | 2.5 | 2.5 | 0 |
| 481 | T | N | 0.7 | -3.5 | -4.2 |
| 482 | P | G | -1.6 | -0.4 | 1.2 |
| 483 | - | V | 0 | 4.2 | 4.2 |
| 484 | P | E | -1.6 | -3.5 | -1.9 |
| 485 | A | G | 1.8 | -0.4 | -2.2 |
| 486 | L | F* | 3.8 | 2.8 | -1 |

| | | | | | |
|---|---|---|---|---|---|
| 487 | N | N* | -3.5 | -3.5 | 0 |
| 488 | C | C | 2.5 | 2.5 | 0 |
| 489 | Y | Y* | -1.3 | -1.3 | 0 |
| 490 | W | F | -0.9 | 2.8 | 3.7 |
| 491 | P | P | 2.8 | 2.8 | 0 |
| 492 | L | L | 3.8 | 3.8 | 0 |
| 493 | N | Q* | -3.5 | -3.5 | 0 |
| 494 | D | S | -3.5 | -0.8 | 2.7 |
| 495 | Y | Y | -1.3 | -1.3 | 0 |
| 496 | G | G* | -0.4 | -0.4 | 0 |
| 497 | F | F | 2.8 | 2.8 | 0 |
| 498 | Y | Q* | -1.3 | -3.5 | -2.2 |
| 499 | T | P | 0.7 | -1.6 | -2.3 |
| 500 | T | T* | -0.7 | -0.7 | 0 |
| 501 | T | N* | 0.7 | -3.5 | -4.2 |
| 502 | G | G* | -0.4 | -0.4 | 0 |
| 503 | I | V | 4.5 | 4.2 | -0.3 |
| 504 | G | G | -0.4 | -0.4 | 0 |
| 505 | Y | Y* | -1.3 | -1.3 | 0 |

| | | | | | |
|---|---|---|---|---|---|
| 506 | Q | Q | -3.5 | -3.5 | 0 |
| 507 | P | P | -1.6 | -1.6 | 0 |
| 508 | Y | Y | -1.3 | -1.3 | 0 |
| 509 | R | R | -4.5 | -4.5 | 0 |
| 510 | V | V | 4.2 | 4.2 | 0 |
| 511 | V | V | 4.2 | 4.2 | 0 |
| 512 | V | V | 4.2 | 4.2 | 0 |
| 513 | L | L | 3.8 | 3.8 | 0 |
| 514 | S | S | -0.8 | -0.8 | 0 |
| 515 | F | F | 2.8 | 2.8 | 0 |
| 516 | E | E | -3.5 | -3.5 | 0 |
| 517 | L | L | 3.8 | 3.8 | 0 |
| 518 | L | L | 3.8 | 3.8 | 0 |
| 519 | N | H | -3.5 | -3.2 | 0.3 |
| 520 | A | A | 1.8 | 1.8 | 0 |
| 521 | P | P | -1.6 | -1.6 | 0 |
| 522 | A | A | 1.8 | 1.8 | 0 |
| 523 | T | T | -0.7 | -0.7 | 0 |

| | | | | | |
|---|---|---|---|---|---|
| 524 | V | V | 4.2 | 4.2 | 0 |
| 525 | C | C | 2.5 | 2.5 | 0 |
| 526 | G | G | -0.4 | -0.4 | 0 |
| 527 | P | P | -1.6 | -1.6 | 0 |
| 528 | K | K | -3.9 | -3.9 | 0 |
| 529 | L | K | 3.8 | -3.9 | -7.7 |
| 530 | S | S | -0.8 | -0.8 | 0 |
| 531 | T | T | -0.7 | -0.7 | 0 |

**Table S2.** Statistical analysis of opening degree at the RBD/hACE-2 interface at different salt concentrations. The opening is quantified based on the angle between two vectors corresponding to the $C_\alpha$ coordinates of hACE-2 α-helix (residues 38-49) and Spike RBD β-strand (492-494 residues), shown in **figure 2f** in the main text.

| NaCl concentration | mean (°) | maximum opening (°) | standard deviation (°) |
|---|---|---|---|
| 150 | 25 | 36 | 3 |
| 500 | 26 | 36 | 3 |
| 1000 | 28 | 41 | 4 |

**Table S3.** Statistical analysis of distances between the centers of masses of the RBD and hACE-2 at different salt concentrations.

| NaCl concentration | mean (Å) | maximum distance (Å) | standard deviation (Å) |
|---|---|---|---|
| 150 | 47.6 | 50.0 | 1.0 |
| 500 | 48.3 | 50.4 | 0.6 |
| 1000 | 49.5 | 52.7 | 0.8 |

**Table S4**. Hydrogen bond occupancy (%) analysis of the RBD residues that make contact with hACE-2 along MD simulations. Table shows intramolecular interactions (RBD (intra)) and intermolecular interactions (hACE-2 (inter)). Cut-off: hydrogen bond distance of 3 Å between hydrogen and nitrogen or oxygen. sc = side chain and bb = backbone. Residues highlighted in green and orange are located in the E1 and E2 regions, respectively.

| **hACE-2/RBD interface residues** | | | **HB occupancy (%)** | | |
|---|---|---|---|---|---|
| **RBD** | **RBD (intra)** | **hACE-2 (inter)** | **150 mM** | **500 mM** | **1,000 mM** |
| **K417** | | D30 | 46 | 46 | 10 |
| | | H34 | 0 | 0 | 15 |
| | Q406 | | 0 | 0 | 12 |
| **G446** | | Q42 | 4 | 4 | 7 |
| | Q498 | | 0 | 0 | 4 |
| **Y449** | | D38 | 75 | 75 | 75 |
| | | Q42 | 0 | 0 | 6 |
| **Y453** | | H34 | 4 | 4 | 0 |
| **N487** | | Y83 | 28 | 28 | 0 |
| | Y489 | | 41 | 42 | 0 |
| | Q474 | | 0 | 0 | 8 |
| | T478 | | 0 | 0 | 10 |
| | T478 | | 0 | 0 | 19 |
| **Y489** | N487 | | 41 | 41 | 0 |
| | | T27 | 36 | 36 | 47 |
| | | T83 | 0 | 0 | 9 |
| **Q493** | S494 | | 32 | 32 | 22 |
| | | E35 | 46 | 46 | 45 |
| | | K31 | 7 | 7 | 9 |
| **Y495** | E406 | | 41 | 41 | 62 |
| **G496** | | K353 | 10 | 10 | 6 |
| **Q498** | N501 | | 24 | 24 | 17 |
| | G446 | | 2 | 2 | 4 |

| | | | | | |
|---|---|---|---|---|---|
| | | K353 | 11 | 11 | 15 |
| | | D38 | 70 | 70 | 62 |
| **T500** | | Y41 | 45 | 45 | 34 |
| **N501** | Q498 sc | | 24 | 24 | 17 |
| | Q498 bb | | 9 | 9 | 10 |

**Table S5.** Covariance analysis of RBD/hACE-2 complex at different NaCl concentrations showing only the intermolecular covariant pairs from MD simulation .

| **150 mM NaCl** | | | | **500 mM NaCl** | | | | **1,000 mM NaCl** | | | |
|---|---|---|---|---|---|---|---|---|---|---|---|
| positive | | negative | | positive | | negative | | positive | | negative | |
| **ACE2** | **RBD** | **ACE2** | **RBD** | **ACE2** | **RBD** | **ACE2** | **RBD** | **ACE2** | **RBD** | **ACE2** | **RBD** |
| K353 | N501 | F555 | F515 | G353 | N501 | | | F28 | F486 | S317 | D364 |
| G354 | N501 | C530 | G431 | | | | | K31 | Y489 | V318 | D364 |
| D355 | N501 | A532 | G431 | | | | | K31 | Q493 | V318 | Y365 |
| K353 | G502 | A533 | G431 | | | | | L79 | G485 | | |
| G354 | G502 | C530 | C432 | | | | | | | | |
| K353 | Y505 | A532 | C432 | | | | | | | | |
| G354 | Y505 | A533 | C432 | | | | | | | | |
| | | C530 | V433 | | | | | | | | |
| | | C530 | C379 | | | | | | | | |
| | | C530 | Y380 | | | | | | | | |
| | | C530 | V382 | | | | | | | | |

**Table S6.** The trajectory of Spike RBD/hACE-2 center-of-mass with the simulation time was fitted to equation $r\,(t) = A + B.e^{-k.t}$, where *k*, *A*, and *B* are the diffusion rate, the amplitude and an offset, respectively.

| NaCl concentration (mM) | A (nm) | B (nm) | *k* ($ns^{-1}$) |
|---|---|---|---|
| 150 | 4.97 ± 0.02 | 0.99 ± 0.15 | 1.18 ± 0.28 |
| 1000 | 5.12 ± 0.02 | 0.77 ± 0.10 | 0.47 ± 0.11 |

*The errors are based on 1σ level of confidence of the probability distribution function.

**Table S7.** Total number of ions detected within 6 Å from each residue at the hACE-2 interface along free diffusion.

| Residue number | number of ions (150 mM) | ions/ns (150 mM) | number of ions (1,000 mM) | ions/ns (1,000 mM) |
|---|---|---|---|---|
| 24 | 0 | 0 | 20 | 1 |
| 27 | 1 | 0.05 | 43 | 2.15 |
| 28 | 0 | 0 | 0 | 0 |
| 30 | 37 | 1.85 | 62 | 3.1 |
| 31 | 0 | 0 | 5 | 0.25 |

| | | | | |
|---|---|---|---|---|
| 34 | 1 | 0.05 | 124 | 6.25 |
| 35 | 76 | 3.8 | 251 | 12.55 |
| 37 | 0 | 0 | 3 | 0.15 |
| 38 | 11 | 0.55 | 86 | 4.3 |
| 41 | 0 | 0 | 0 | 0 |
| 42 | 0 | 0 | 109 | 5.45 |
| 45 | 0 | 0 | 6 | 0.3 |
| 82 | 56 | 2.8 | 347 | 17.35 |
| 83 | 17 | 0.85 | 101 | 5.05 |
| 330 | 23 | 1.15 | 622 | 31.1 |
| 353 | 0 | 0 | 99 | 4.95 |
| 354 | 0 | 0 | 162 | 8.1 |
| 355 | 0 | 0 | 51 | 2.55 |
| 357 | 0 | 0 | 0 | 0 |
| 393 | 340 | 17 | 193 | 9.65 |

**Table S8.** Total number of ions detected within 6 Å from each residue at the RBD interface along the free diffusion experiment. Residues colored in green belong to the E1 region, while those colored orange belong to the E2 region.

| RBD residue | number of ions (150 mM) | ions/ns (150 mM) | number of ions (1,000 mM) | ions/ns (1,000 mM) |
|---|---|---|---|---|
| 417 | 73 | 3.65 | 159 | 7.95 |
| 446 | 26 | 1.3 | 661 | 33.05 |
| 453 | 0 | 0 | 0 | 0 |
| 455 | 52 | 2.6 | 0 | 0 |
| 456 | 1 | 0.05 | 70 | 3.5 |
| 475 | 4 | 0.2 | 102 | 5.1 |
| 486 | 43 | 2.15 | 168 | 8.4 |
| 487 | 0 | 0 | 38 | 1.9 |
| 489 | 23 | 1.15 | 58 | 2.9 |
| 493 | 86 | 4.3 | 172 | 8.6 |
| 495 | 0 | 0 | 34 | 1.7 |
| 496 | 0 | 0 | 72 | 3.6 |
| 498 | 0 | 0 | 12 | 0.6 |
| 500 | 3 | 0.15 | 412 | 20.6 |
| 501 | 0 | 0 | 193 | 9.65 |
| 502 | 3 | 0.15 | 331 | 16.55 |

| | | | | |
|---|---|---|---|---|
| 505 | 0 | 0 | 2 | 0.1 |

**Table S9.** Binding kinetics obtained by surface plasmon resonance experiments. Two constructs of purified recombinant human ACE-2 (residues from 18 to 740) were used: hACE-2 with C-terminal Fc tag and Biotinylated hACE-2 with C-terminal polyhistidine and avidin tags. Purified hACE-2 constructs were covalently immobilized at the surface of the sensor chip, and different concentrations of SARS-CoV-2 recombinant purified proteins were applied. Four different spike protein constructs were used, the RBD (residues 319 to 537), the S1 subunit (residues 16 to 685), monomeric form of the spike protein (residues 16 to 1213, with mutations at arginines 683 and 685 to alanines: R683A, R685A), and trimeric form of spike protein (residues 16 to 1213, with mutations at arginines 683 and 685 to alanines: R683A, R685A). Shown data are representative of three independent experiments.

| **Dimer hACE-2$_{18\text{-}740}$** | **Immobilized Level (RU)** | **Spike constructs** | **NaCl (mM)** | $k_a$ **(M$^{-1}$s$^{-1}$)** | $k_d$ **(s$^{-1}$)** | $K_D$ **(M)*** | **Rmax (RU)** | **Chi² (RU²)** |
|---|---|---|---|---|---|---|---|---|
| ACE2, Fc tag (110 kDa) | 156.2 | RBD$_{319\text{-}537}$ (26.5 kDa) | 50 | 264 $10^4$ | 11.4 $10^{-3}$ | **4.3 $10^{-9}$** | 63.9 | 1.5 |
| | 163.8 | | 150 | 341 $10^4$ | 3.7 $10^{-3}$ | **1.1 $10^{-9}$** | 44.4 | 0.9 |
| | 160.8 | | 500 | 97.1 $10^4$ | 7.5 $10^{-3}$ | **7.8 $10^{-9}$** | 34.6 | 0.2 |

| | | | | | | | | |
|---|---|---|---|---|---|---|---|---|
| | 159.7 | | 1000 | $44.8\ 10^{4}$ | $3.6\ 10^{-3}$ | **$8.0\ 10^{-9}$** | 30.8 | 0.2 |
| ACE2, Fc tag (110 kDa) | 166.3 | Trimer $S_{16\text{-}1213}$ (R683A, R685A) (138 kDa) | 50 | $4.3\ 10^{4}$ | $0.8\ 10^{-3}$ | **$17.4\ 10^{-9}$** | 209.3 | 7.0 |
| | 208.5 | | 150 | $4.1\ 10^{4}$ | $0.7\ 10^{-3}$ | **$16.2\ 10^{-9}$** | 165.4 | 1.5 |
| | 178.3 | | 500 | $2.6\ 10^{4}$ | $0.2\ 10^{-3}$ | **$8.0\ 10^{-9}$** | 176.9 | 0.2 |
| | 156.8 | | 1000 | $1.2\ 10^{4}$ | $0.4\ 10^{-3}$ | **$3.3\ 10^{-9}$** | 176.4 | 0.1 |
| ACE2, Fc tag (110 kDa) | 164.8 | $S1_{16\text{-}685}$ subunit (76.9 kDa) | 150 | $12.4\ 10^{4}$ | $1.5\ 10^{-3}$ | **$12.3\ 10^{-9}$** | 111.1 | 0.8 |
| | 158.5 | | 1000 | $8.3\ 10^{4}$ | $1.1\ 10^{-3}$ | **$13.3\ 10^{-9}$** | 100.6 | 3.2 |
| ACE2, Avi Tag (87.2 kDa) | 114.1 | Monomer $S_{16\text{-}1213}$ (R683A, R685A) (134.6 kDa) | 285 | $7.3\ 10^{4}$ | $0.9\ 10^{-3}$ | **$12.3\ 10^{-9}$** | 167.8 | 1.6 |
| | 103.2 | | 500 | $5.7\ 10^{4}$ | $1.0\ 10^{-3}$ | **$17.8\ 10^{-9}$** | 206.1 | 1.5 |
| | 99.6 | | 1000 | $4.5\ 10^{4}$ | $0.7\ 10^{-3}$ | **$15.8\ 10^{-9}$** | 242.4 | 0.7 |

* $K_D = k_d/k_a$
